# Multiomics analysis reveals extensive epigenome remodeling during cortical development

**DOI:** 10.1101/2020.08.07.241828

**Authors:** Florian Noack, Silvia Vangelisti, Madalena Carido, Faye Chong, Boyan Bonev

## Abstract

Despite huge advances in stem-cell, single-cell and epigenetic technologies, the precise molecular mechanisms that determine lineage specification remain largely unknown. Applying an integrative multiomics approach, e.g. combining single-cell RNA-seq, single-cell ATAC-seq together with cell-type-specific DNA methylation and 3D genome measurements, we systematically map the regulatory landscape in the mouse neocortex *in vivo*. Our analysis identifies thousands of novel enhancer-gene pairs associated with dynamic changes in chromatin accessibility and gene expression along the differentiation trajectory. Crucially, we provide evidence that epigenetic remodeling generally precedes transcriptional activation, yet true priming appears limited to a subset of lineage-determining enhancers. Notably, we reveal considerable heterogeneity in both contact strength and dynamics of the generally cell-type-specific enhancer-promoter contacts. Finally, our work suggests a so far unrecognized function of several key transcription factors which act as putative “molecular bridges” and facilitate the dynamic reorganization of the chromatin landscape accompanying lineage specification in the brain.

## Introduction

Cellular identity is established by the complex interplay between transcriptional regulators, cis-regulatory elements and chromatin landscape, which takes place within the physical constraints imposed by the 3D nuclear architecture (Schoenfelder and Fraser, 2019). These different layers of molecular interactions form the basis of Gene Regulatory Networks (GRNs), which ensure the precise temporal and spatial regulation of gene expression. Although our understanding of the molecular cascades involved in gene regulation has grown considerably over the last couple of years, the exact multi-layered mechanisms leading to lineage specification or developmental plasticity remain to be understood.

Until very recently, the epigenetic and molecular programs that govern lineage commitments have only been studied *in vitro*, mostly in bulk and restricted to only a few regulatory layers (Rubin et al., 2017; Ziller et al., 2015). Most tissues however, consist of heterogeneous cell populations which can be generally associated to distinct stages of lineage commitment. Single-cell technologies provide access to the dynamics of gene expression and chromatin accessibility, even in complex tissues and *in vivo* (Granja et al., 2019; Telley et al., 2019). Despite the tremendous potential of these technological advances to unravel the dynamics of lineage commitment, studies that integrate single-cell data with DNA methylation- and 3D genome organization patterns remain scarce (Argelaguet et al., 2019; Lee et al., 2019).

Many of these regulatory layers converge on distal enhancers, which represent the key building blocks of GRNs in eukaryotes. Changes in histone modifications/accessibility, as well as the binding of cell-type-specific transcription factors (TFs) have been proposed to explain the relationship between enhancer activation and gene expression (Gorkin et al., 2020; He et al., 2020). Although enhancer-promoter contacts have been previously shown to be highly dynamic (Bonev et al., 2017; Javierre et al., 2016), many examples of pre-looped architecture have been identified (Ghavi-Helm et al., 2019) and the causal relationships between chromatin looping, epigenetic signature and tissue-specific gene expression remain to be understood.

The mammalian embryonic neocortex represents a unique system to study the mechanisms of GRNs and enhancer dynamics *in vivo*. It is mostly composed of neural stem cells (NSC), intermediate progenitors (IPC) and postmitotic neurons in various stages of neuronal maturation (Götz and Huttner, 2005; Govindan and Jabaudon, 2017). Both chromatin organization (Preissl et al., 2018; Telley et al., 2019) as well as epigenetic regulation (Pereira et al., 2010) have been implicated in in determining fate choices in the cortex from studies on transcriptional dynamics in bulk (Aprea et al., 2013) and at the single-cell level (Loo et al., 2019; Telley et al., 2016, 2019).

To assess how chromatin dynamics across multiple layers govern GRNs in the developing neocortex *in vivo*, here we integrate, for the first time, single-cell transcriptomic and epigenomic data with cell-type-specific bulk DNA methylation and 3D genome organization using the mouse neocortex as a model system. We identify key transcription factors to be associated with extensive changes in chromatin accessibility and provide evidence that chromatin remodeling occurs predominantly at distal regulatory elements. Furthermore, we identify thousands of novel cell-type-specific enhancer-gene pairs and show that although enhancer activation appears to precede gene expression, only a subset of enhancers acts as truly lineage-priming. Using an improved method to simultaneously map DNA methylation and 3D genome architecture of immunoFAC-sorted cells, we show that although regulatory contacts between enhancers and promoters are in general correlated with gene expression, considerable heterogeneity in both contact strength and dynamics exists. Finally, we identify a new role for transcription factors as key components of the cell-type-specific reorganization of the chromatin landscape during lineage specification in the mouse brain.

## Results

### Profiling gene expression dynamics during neuronal differentiation at single-cell resolution

To identify changes in the gene regulatory landscape upon differentiation at the single-cell level, we decided to construct independent transcriptomic and epigenetic maps of cortical development *in vivo*. To this end, we dissected the somatosensory area of the E14.5 neocortex and performed in parallel scRNA-seq and scATAC-seq (10x Genomics; Methods) in biological duplicates (Figure 1A).

**Figure 1.**
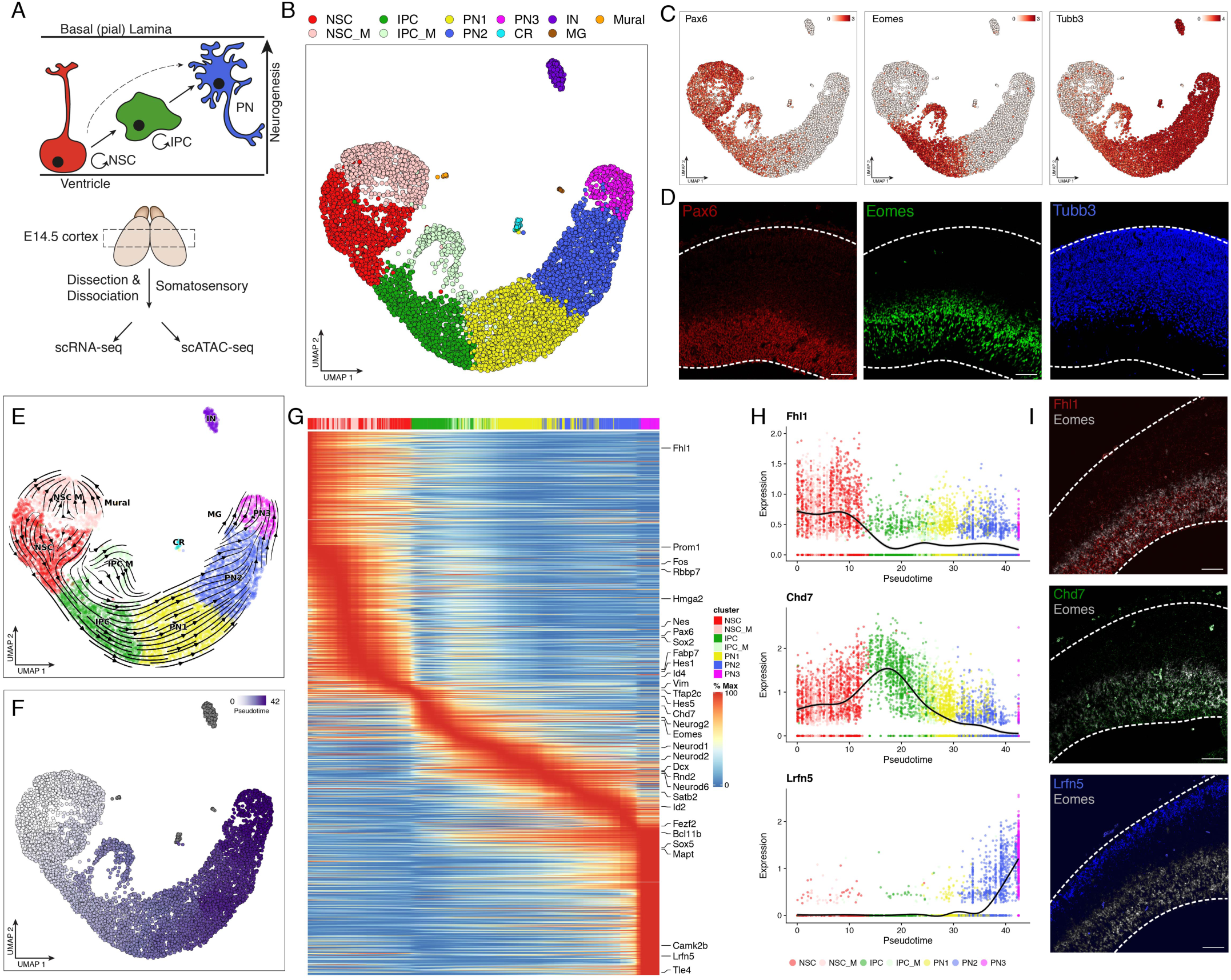
scRNA-seq analysis of mouse E14.5 cortical development. (A) Schematic representation of the model system and the experimental approach. Neural stem cells (NSC), intermediate progenitor cells (IPC) and projection neurons (PN). (B) scRNA-seq UMAP projection of 7469 cells derived from the E14.5 somatosensory cortex. IN= interneurons, CR= Cajal-Retzius neurons, MG= Microglia, Mural= Mural cells, _M indicated the corresponding mitotic (G_2_-M) population. (C) UMAP visualization with expression levels of the indicated marker genes (D) Representative immunofluorescence images of the indicated genes in coronal sections of a E14.5 cortex. Scale = 50µm. (E) Direction of neuronal differentiation inferred from estimated RNA velocities and plotted as streamlines on the UMAP. (F) Trajectory analysis depicting the inferred pseudotime on the UMAP projections. (G) Pseudotime heatmap ordering of the top 3000 most variable genes across neural differentiation. (H) Expression levels of the indicated genes across differentiation. Each dot shows the expression in an individual pseudotime-ordered cell, the line represents the smoothed fit of expression levels. (I) Representative FISH images of the depicted genes in coronal sections of a E14.5 cortex. Scale = 50µm.

To gain insight into the transcriptional dynamics during neuronal differentiation, we first examined our scRNA-seq data. The two replicates were highly comparable (Figure S1A; *r*=0.979) and we obtained 7469 cells in total that passed quality control, with a median of 16812 UMIs and 4439 genes/per cell (Figure S1B-F). Next, we clustered the cells using a graph-based approach (Stuart et al., 2019), annotated the resulting clusters based on known marker genes (Figure 1B and S1G). The resulting 11 clusters were highly reproducible between biological replicates (Figure S1H-I) and captured all major cell types of the developing cortex: neural stem cells (NSC), intermediate progenitor cells (IPC) and projection neurons (PN1-3), as well as rarer and transcriptional distinct cell types. We also confirmed the expression of selected marker genes using immunohistochemistry (Figure 1D).

The observed UMAP projection displayed a gradual progression from NSC to IPC and then to PN (Figure 1B), reflecting a continuum of transcriptional changes compared to the rather distinct cell types reported in the adult cortex (Zeisel et al., 2018). Next, we used two independent methods to infer the developmental trajectory and identify changes in gene expression: a generalized RNA velocity approach (Figure 1E, S1J; Bergen et al., 2020) and Monocle3 (Figure 1F-G, S1K; Cao et al., 2019). Both approaches were highly comparable, suggesting a transition from NSC into IPC, followed by neuronal commitment and maturation (PN1-3). Consistent with a previous lineage-tracing results (Telley et al., 2016), our trajectory analysis revealed transcriptional waves of key neurogenic factors in NSC (Hes1, Id4, Hes5), IPC (Neurog2, Eomes), PN1 (Neurod2), PN2 (Rnd2) and PN3 (Mapt) (Figure S1L-M), Besides these known factors, we identified a number of novel genes exhibiting cell-type-specific expressions along the studied differentiation trajectory, and which we could verify by independent FISH-experiments (Figure 1G-I). For example Fhl1, a skeletal and cardiac muscle protein involved in muscular dystrophy (Cowling et al., 2011) was exclusively expressed in NSC. Chd7, a member of the chromodomain family of chromatin remodellers and associated with the CHARGE syndrome (Vissers et al., 2004), was also expressed in NSC but became upregulated in IPC, consistent with previous reports in the adult brain (Feng et al., 2013) and the developing cerebellum (Feng et al., 2017). Finally, the cell adhesion protein Lrfn5, which has been linked to autism (de Bruijn et al., 2010), developmental delays and mental retardation (Mikhail et al., 2011) was upregulated upon neuronal maturation where it is most likely involved in synaptic formation (Choi et al., 2016).

Collectively, our scRNA-seq data consistently recapitulates known transcriptional dynamics during neurogenesis, reveals cell-type-specific expression of novel genes and provides a molecular roadmap to investigate the influence of epigenetic regulation on gene expression in the context of corticogenesis.

### Accessibility changes at distal regulatory regions and TF binding motifs identified using scATAC-seq

In order to dissect the molecular mechanism orchestrating the observed transcriptional dynamics, we next focused on the scATAC-seq analysis. Again, the two biological replicates were highly correlated (Figure S2A; *r*=0.975), revealed a very high number of unique fragments per cell and showed strong enrichment at the transcription start site (TSS) (Figure S2B). After filtering for low-quality cells and potential doublets we acquired in total 5877 cells with a median of 46899 fragments per cell and a characteristic fragment size distribution and periodicity (Figure S2C-E), making this, to our knowledge, one of the highest resolution scATAC-seq datasets available.

First, we counted the number of Tn5 insertions in 5 kb genomic windows to generate a high-quality union peak set with a fixed length of 501 bp, as previously described (Granja et al., 2019). Next, we determined the number of fragments per peak, followed by dimensionality reduction using latent semantic indexing (LSI), batch correction using Harmony (Korsunsky et al., 2019), graph-based clustering and UMAP-visualization. We identified 7 highly reproducible clusters (Figure 2A, S2F-G), which we subsequently annotated based on the gene body accessibility of known marker genes (Figure S2H-I). Similar to the scRNA-seq, we observed again that NSC, IPC and projection neurons (PN1, PN2) do not form separate clusters but were rather associated with a gradual progression in cell state (Figure 2A). However no distinct mitotic clusters of the progenitor cell types were identified, presumably due to the high similarities of the chromatin accessibility landscape during different cell cycle phases (Estève et al., 2020; Hsiung et al., 2015; Ma et al., 2020). Studying the pro-neuronal gene Dll1 locus as an example, we were able to observe chromatin accessibility dynamics at the single-cell level, and particularly at distal regions (Figure 2B).

**Figure 2.**
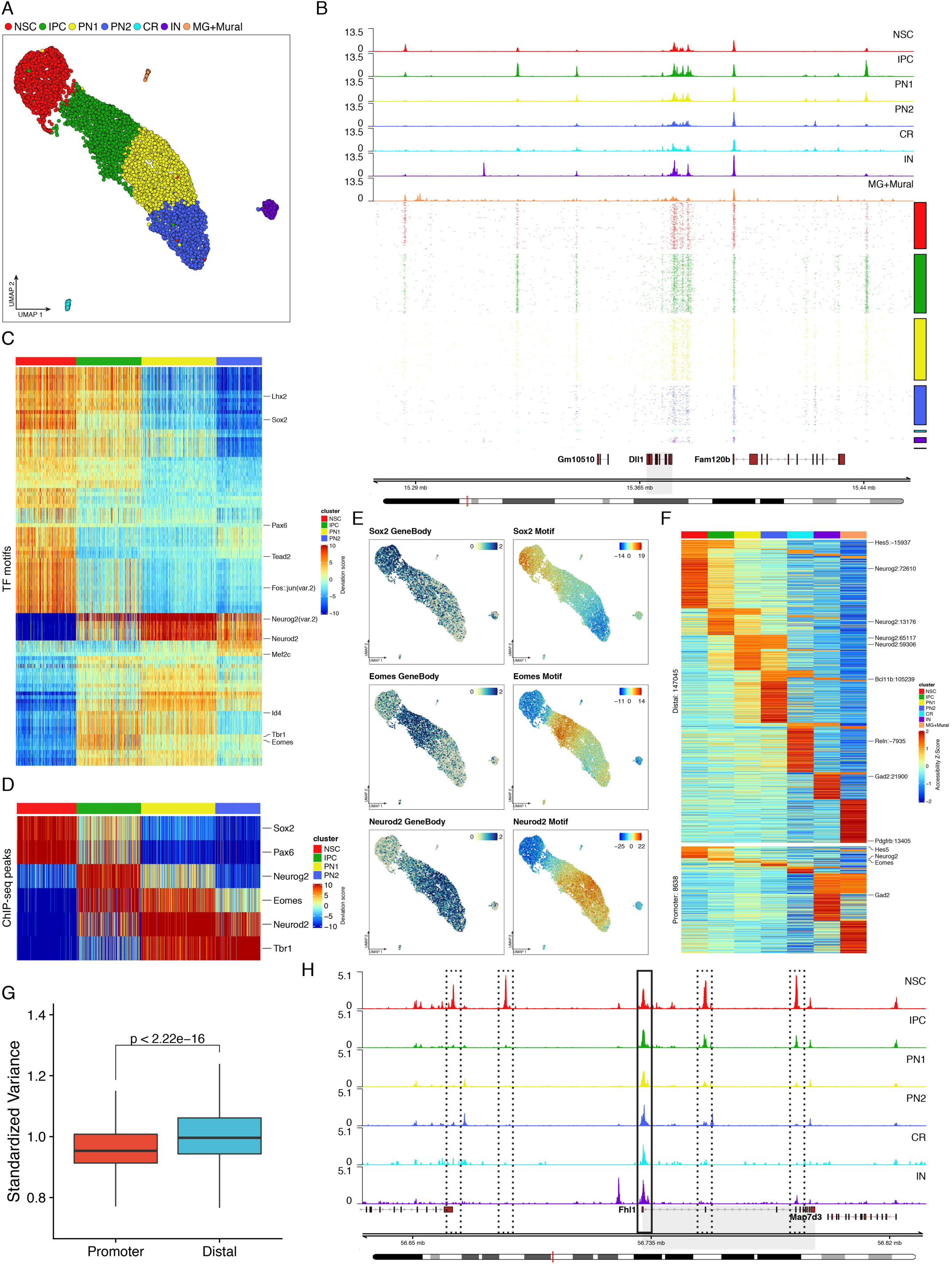
scATAC-seq identifies dynamic TF motifs and variable distal regulatory elements. (A) scATAC-seq UMAP projection of 5877 cells derived from the E14.5 somatosensory cortex. (B) Genomic tracks showing the accessibility of aggregated scATAC-seq clusters (top) and of 1000 random single-cells (bottom) at the *Dll1* gene locus. (C-D) Heatmap clustering of chromVAR bias-corrected accessibility deviations for the most 100 variably expressed TFs (C) or based on ChIP-seq peaks (D). Single-cell cluster identities are indicated on top of the plot. (E) UMAP projections, colored by gene body accessibility of the indicated TFs or by their chromVAR motif bias-corrected deviations (F) Heatmap of aggregated accessibility Z-scores (per cluster) of differentially accessible peaks in distal regions (≥± 5kb from TSS, top) or within promoters (≤± 500bp from TSS, bottom). Labels indicate the name and the distance in bp to the nearest TSS. (G) Box-whisker plot, showing the standardized variance between differential peaks within promoter or distal regions. Statistical significance is calculated using wilcoxon rank-sum test. (H) Genomic tracks showing the accessibility of aggregated scATAC-seq clusters at the Fhl1 gene locus (highlighted in grey). The promoter region (black rectangle) and four putative regulatory elements (dashed black rectangle) are also highlighted.

To identify key transcription factors whose binding correlates with changes in chromatin accessibility, we used ChromVAR (Figure 2C; Fornes et al., 2020; Schep et al., 2017). TFs associated with a high motif accessibility in NSC included classical neural TFs such as Sox2, Pax6 or Lhx2, as well as Tead2 and c-Fos/Jun (AP-1), which have been recently described as important in NSC (Mukhtar et al., 2020; Pagin et al., 2020). Conversely, neurogenic transcription factors such as Eomes and Neurod2 were associated with a strong increase in motif accessibility during the transition from NSC to IPC, and IPC to PN1 respectively (Figure 2C and 2E). Since TF-binding motifs are only predictive for real binding events, we verified our findings using publicly available ChIP-seq data (Schep et al., 2017). Overall, both motif- and peak-based accessibility showed a high correlation with the expression level of the respective transcription factors (Figure 2C-E and Figure 1).

To determine how accessibility at regulatory elements changes upon differentiation, we first identified differentially accessible regions as previously described (Granja et al., 2019) and grouped them subsequently into distal and promoter associated elements (Figure 2F). The majority of changes in accessibility occurred within distal elements rather than on promoters, indicating that the former are more cell-type-specific than the latter (Figure 2F-G and S2J). To corroborate this finding, we visualized the Fhl1 locus (expressed only in NSC – Figure 1). Parallel to Fhl1 downregulation, four distal elements became less accessible, while the promoter region remained constant (Figure 2H). These findings are consistent with the hypothesis that promoter accessibility, while required, may not be sufficient for transcription (Corces et al., 2016; Klemm et al., 2019).

Collectively, our scATAC-seq data identify major reorganization of the chromatin landscape during neural differentiation at the single-cell level. This extensive remodelling is associated with unique binding patterns of cell-type-specific transcription factors. We also demonstrate that the accessibility at distal regulatory elements (such as enhancers) is highly dynamic and frequently correlated with changes in gene expression, compared to relatively invariant/static promoters.

### Lineage-specific enhancer-gene pairs underlie the reorganization of chromatin landscape

To uncover how dynamic accessibility at distal regulatory elements relates to changes in gene expression genome-wide, we employed an approach pioneered by the Greenleaf Lab (Granja et al., 2019, 2020). First, we integrated the scRNA-seq and scATAC-seq data by using canonical correlation analyses (Stuart et al., 2019), which resulted in overall high prediction scores, minimal cross-annotations between the datasets and high intermixed projection of the integrated cells on a joint UMAP (Figure S3A-D). Next, we calculated the pairwise correlation between changes in gene expression and chromatin accessibility of nearby distal peaks (±500 kb, at least 5 kb away from a TSS) to link potential distal regulatory elements and their putative target genes. These analyses identified 16978 positively correlated pairs (r≥0.35, FDR≤0.1) with each gene being connected to a median of 3 distal regions (Figure S3E). Positively correlated distal regions were usually located closer to their predicted target genes (Figure S3F) and were characterized by overall higher accessibility compared to non-correlated pairs (also referred to as control pairs, -0.35≥r≤0.35, FDR>0.1) but not compared to negatively correlated pairs (r≤-0.35, FDR≤0.1; Figure S3G).

To address if the identified pairs are cell-type-specific, we clustered the identified distal regions based on their pseudobulk accessibility. Intriguingly, we found that the majority of the newly identified positively correlated pairs (which we will refer to as enhancer-gene pairs for simplicity) were characterized by high cell-type specificity in both enhancer accessibility and target gene expression (Figure 3A). This is further emphasized by the striking contrast to the negatively correlated and control pairs (Figure S3H-I). Finally, we found a significant overlap of our positively (90/334, *p*<6.636e-24) correlated enhancer-gene pairs with validated forebrain enhancers (Visel et al., 2013).

**Figure 3.**
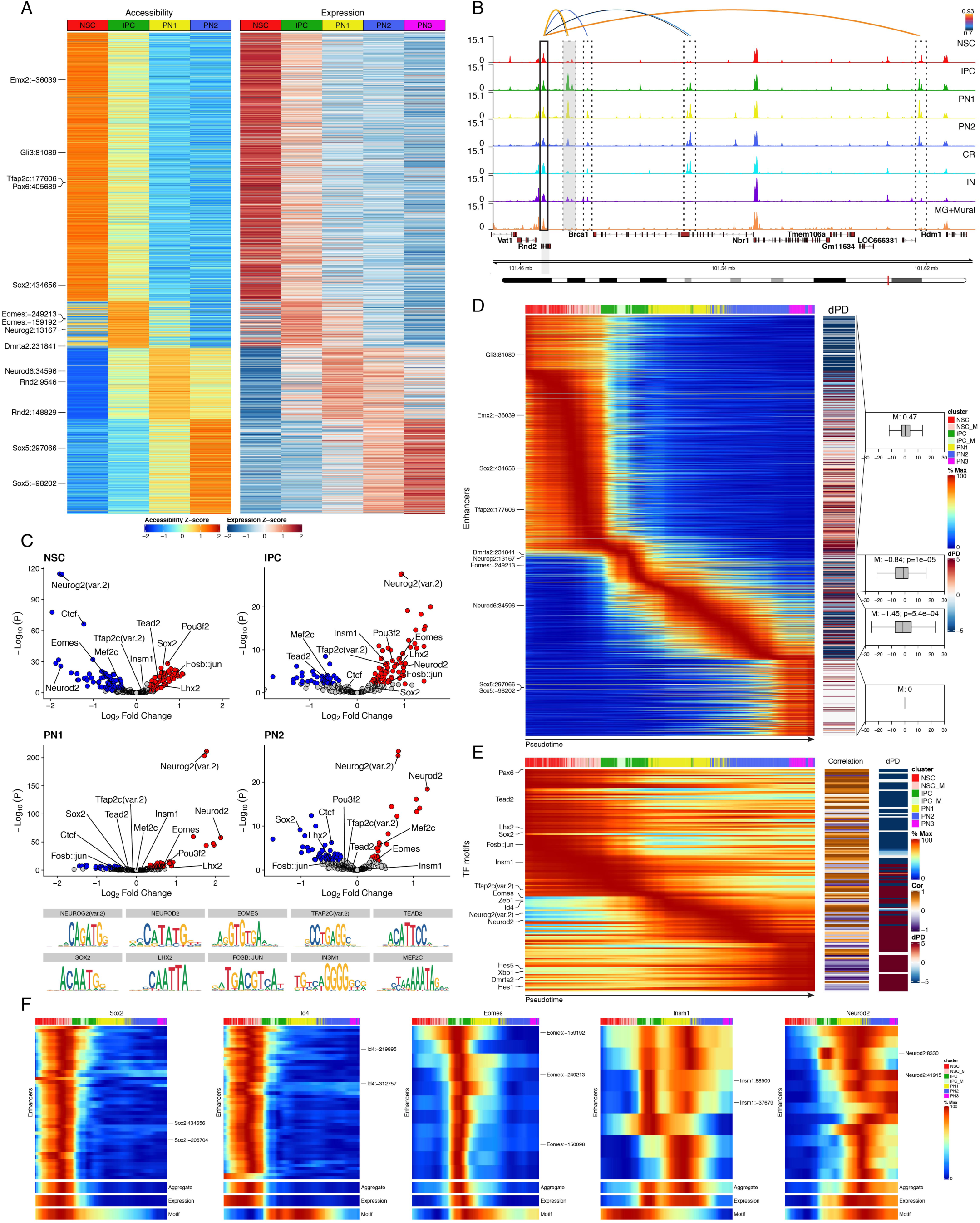
Lineage dynamics of enhancer-gene pairs and transcription factor motifs. (A)Heatmaps of aggregated accessibility of putative enhancers (left) and gene expression levels of their linked genes (right) for each of the 16978 positively correlated enhancer-gene pairs identified (rows). Rows were clustered by enhancer accessibility using feature binarization (Methods). (B) Genomic tracks depicting aggregated accessibility (per cluster) at the Rnd2 gene locus. Arcs represent Rnd2 linked enhancers and their correlation score. The promoter region (black rectangle), previously characterized enhancers (Heng et al., 2008; grey dashed rectangle) and putative predicted enhancers (black dashed rectangle) are highlighted. (C) Scatter plot showing the enrichment of transcription factor binding motifs within cluster-specific positive correlated enhancer-gene pairs. Red and blue dots indicate significantly (log_10_(P) ≥ 2; abs(logFC) ≥ 0.25) enriched or depleted motifs, respectively. The binding motifs of the highlighted TFs are depicted in the bottom. (D) Left, heatmap depicting scaled enhancer accessibility of positive correlated enhancer-gene pairs in individual cells ordered along the integrated pseudotime. Right, heatmap depicting the difference between the pseudo-temporal maxima of enhancer accessibility and expression of the linked gene (referred to as “dPD”) and box-whisker plots with the median of these differences (M). Negative values mean that accessibility precedes gene expression. Significance was calculated using one-sample Wilcoxon signed rank test. (E) Analogous to (D) but heatmaps depict the pseudotemporal ordering of chromVAR scaled deviation scores (left), the correlation between motif accessibility and TF expression (middle) and difference between the pseudo-temporal maxima of motif accessibility and expression of the linked gene (referred to as “dPD”). (F) Pseudotime heatmap ordering of enhancer accessibility (individual or aggregate), linked gene expression and motif accessibility of indicated TFs.

To further validate these findings, we focused on the Rnd2 locus, a gene involved in neuronal migration (Heng et al., 2008), which was up-regulated in IPC and mainly expressed in newborn neurons (PN1) (Figure S1L-M). As observed before, we found no correlation between Rnd2 expression and its promoter accessibility but we identified several distal regions, which correlated positively with Rnd2 expression (Figure 3B). Some of those included previously validated Rnd2 enhancers (Heng et al., 2008) but others were completely novel. Importantly, the closest genes of these putative new Rnd2 enhancers were either not expressed (Gm11634, LOC666331) or their expression pattern was not correlated to the enhancer accessibility (Brca1, Nbr1, Rdm1) (Figure 3B).

To address which TFs are associated with the identified cell-type-specific enhancers, we calculated the motif enrichment for TFs expressed during neurogenesis (Figure 3C). We found that some NSC-specific TFs such as Sox2 and Tead2 were enriched only in NSC-specific enhancers, while others such as Lhx2 and Pou3f2 (also known as Brn2) were enriched in both NSC and IPC enhancers. Conversely, neurogenic TFs such as Neurog2, Eomes and Neurod2 were depleted in NSC enhancers but became strongly enriched in both IPC and PN1-2 specific enhancers. Importantly, this approach allowed us to identify Insm1 (Tavano et al., 2018) and Tfap2c (Pinto et al., 2009), two TFs with a described role in inducing and regulating IPC fate, to be specifically enriched within IPC-specific enhancer-gene pairs. Interestingly, Ctcf – a transcription factor frequently associated with 3D genome architecture (Bonev and Cavalli, 2016), was actually depleted in almost all clusters.

To determine the temporal relationship between enhancer accessibility and gene expression, we ordered the positively correlated pairs based on their enhancer accessibility as a function of the integrated pseudotime (Figure 3D). This systematic analysis showed that for the transient cell states (IPC, PN1 and PN2) enhancer accessibility significantly precedes upregulation of their linked gene despite overall strong correlation (Figure3A and Methods). Similar TF motif-centric analysis (Figure 3E) showed that motif accessibility and expression for transcriptional activators such as Pax6, Sox2, Eomes or Neurog2 were highly correlated, while known repressors such as Insm1, Id4 and Hes1 displayed a strong negative correlation. Furthermore, we observed that expression precedes motif accessibility for the majority of transiently expressed TFs.

To address if all predicted enhancers of a gene are characterized by similar accessibility dynamics, we visualized the changes in accessibility of individual enhancers linked to a transcription factor, its expression and the average accessibility of its binding motifs (Figure 3F, S3J). While the majority of the linked enhancers became accessible shortly prior or at the onset of gene expression, some enhancers (such as Eomes:-159192 and Neurod2:8330) acquired accessibility considerably earlier, suggesting that they may act in lineage-priming (Figure 3F) and could be bound by different TFs (Figure S3K).

In summary, the analysis of the integrated scRNA-seq and scATAC-seq datasets allowed us to identify matched enhancer-gene pairs, which led to several novel findings. First, we find both known and novel TFs particularly enriched in the cell-type-specific enhancers. Second, we provide additional support for a long-standing hypothesis in the field, namely that enhancer activation precedes gene expression, although only a subset of enhancers appears to be truly lineage-priming. Finally, the identification of enhancer-gene pairs in single cells, in vivo sets the stage to systematically interrogate how TFs facilitate and maintain GRNs during development.

### Enhancer-gene pairs reveal target genes of neurogenic transcription factors

Identifying the direct downstream targets of TFs is one of the most challenging steps in reconstructing GRNs. Although best results are generally achieved by combining ChIP-seq and misexpression analysis (Berest et al., 2019; Feng et al., 2020), such data remain mostly unavailable. We developed an approach using the obtained enhancer-gene pairs in order to predict bona-fide target genes of (cell-type) specific transcription factors (Methods). We focused first on Eomes, a transcriptional activator expressed in IPC, known to support neuronal expansion (Figure 1C-D; Sessa et al., 2008). Eomes motif was enriched specifically in IPC enhancer-gene pairs (Figure 3C) and displayed the highest accessibility within the IPC cluster (Figure 4A). We verified the putative downstream targets of Eomes (Figure 4B) using three orthogonal approaches. First, gene ontology (GO) analysis identified categories, reminiscent of the known function of Eomes in the developing brain (Figure 4C; reviewed by Hevner, 2019). Second, an analogous approach using Eomes ChIP-seq peaks instead of binding motifs (Sessa et al., 2017) was characterized by comparable accessibility pattern (Figure S4A), identified very similar target genes (Figure S4B) and resulted in highly overlapping GO terms (Figure S4C). Finally, several putative Eomes targets were previously identified to be either bound by it (Cdh8, Satb2, Dll1) or downregulated upon its ablation (Eomes, Dach1, Sstr2, Azi2, Cdh8, Satb2) (Sessa et al., 2017). Importantly, our approach allowed us to also predict novel direct targets of Eomes, which could not be identified with conventional promoter-based approaches. A representative example is Sstr2, an important receptor for neuronal migration and axon outgrowth (Stumm et al., 2004) (Figure 4D-E).

**Figure 4.**
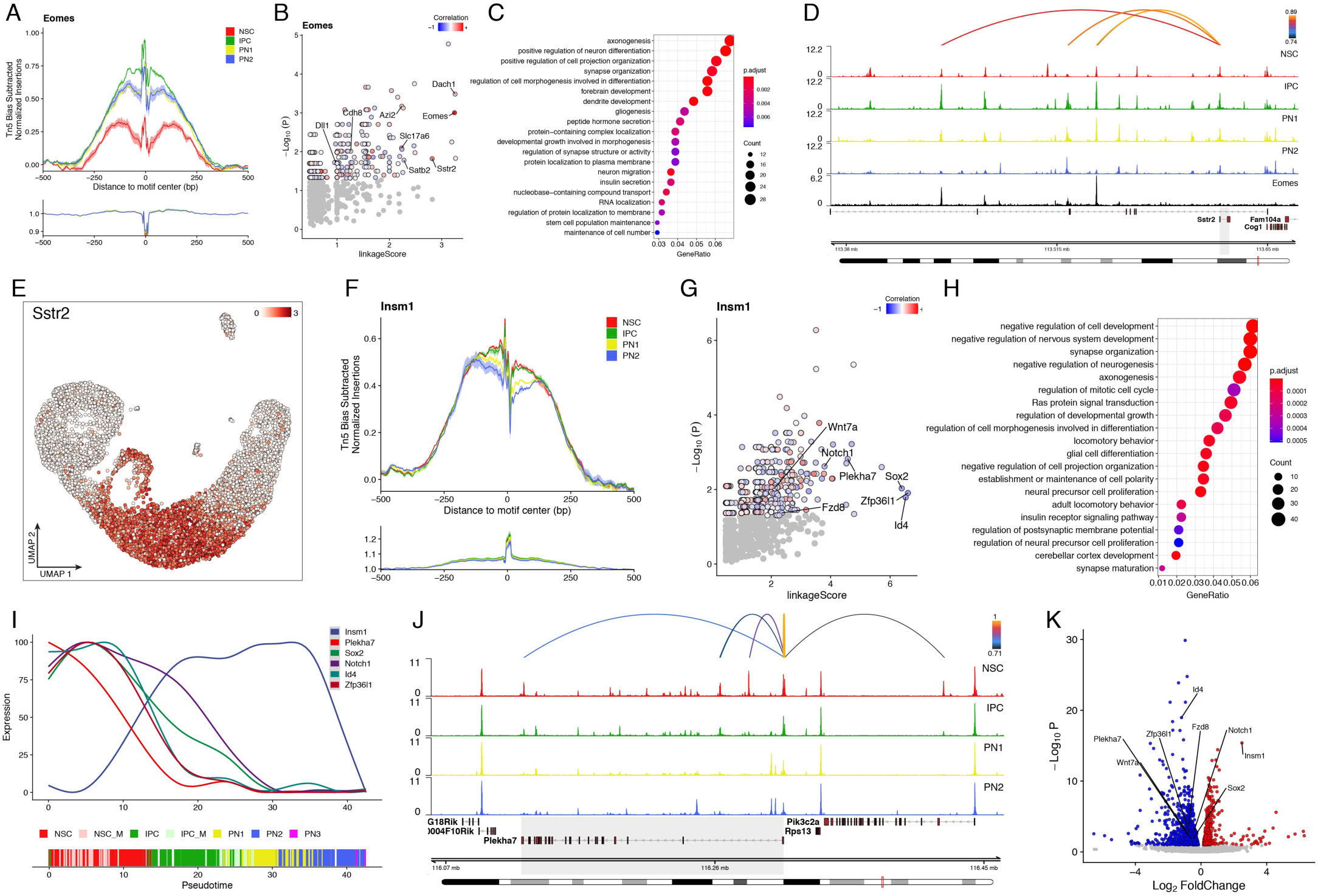
Gene regulatory networks associated with the TF Eomes and Insm1. (A) TF footprint of Eomes binding motif in the indicated scATAC clusters. The Tn5 insertion bias track is shown below. Footprint is calculated using ArchR (Granja et al., 2020) (B) Scatter plot depicting predicted downstream targets of Eomes based on the aggregated correlation between gene expression and accessibility of linked enhancers containing the Eomes motif (linkageScore) versus the statistical significance of the motif enrichment (hypergeometric test). Significant genes are colored based on the Pearson correlation between their expression and the expression of Eomes. (C) Bar plot depicting the GO enriched terms, enrichment scores and gene ratios of the predicted Eomes targets. (D) Genomic tracks depicting aggregated accessibility (per cluster) and Eomes Chip-seq profile (black) at the Sstr2 gene locus. Arcs on top represent linked Sstr2 enhancers, overlapping with Eomes motif and colored by Pearson correlation of the enhancer accessibility and Sstr2 expression. Note the overlap of Eomes binding sites and linked enhancers. (E) scRNA UMAP projection, colored by the expression levels of Sstr2. (F) TF footprint of Insm1 binding motif in the indicated scATAC clusters. (G) as in (B) but based on Insm1 instead of Eomes. (H) Bar plot depicting the GO enriched terms, enrichment scores and gene ratios of the predicted Insm1 targets. (I) Smoothed fit line plots depicting the scaled gene expression levels of Insm1 and some of its predicted downstream target genes across pseudotime. (J) Genomic tracks depicting aggregated accessibility (per cluster) at the known Isnm1 target Plekha7 (highlighted in grey; Tavano et al., 2018). Arcs on top represent linked Plekha7 enhancers, overlapping with Insm1 motif and colored by Pearson correlation of the enhancer accessibility and Plekha7 expression. (K) Volcano plot of significantly (FDR< 0.1) down (blue) and upregulated (red) genes upon overexpression of Insm1 in E14.5 cortex (Tavano et al., 2018).

After verifying our strategy with Eomes, we next asked whether our approach would similarly capture targets of putative repressors. Insm1, has a known function as transcriptional repressor in the NSC to IPC cell differentiation (Farkas et al., 2008; Monaghan et al., 2017; Tavano et al., 2018), was strongly upregulated in IPC (Figure 3F, S3J) and displayed a specific enrichment of its motifs only within IPC enhancer-gene pairs (Figure 3C). While no Insm1 cortex-specific ChIP-seq data exists, we found that Insm1 binding motifs lose accessibility upon differentiation (PN1-2) (Figure 4F). We identified 713 potential target genes (Figure 4G), the majority of which displayed a negative correlation with Insm1 expression (430/713). Overall the identified targets were associated with GO terms such as negative regulation of neurogenesis, fully consistent with Insm1’s repressor function in IPC differentiation (Farkas et al., 2008; Tavano et al., 2018) (Figure 4H). Putative targets included NSC-specific genes such as Sox2, Notch1, Id4, Zfp36l1 and the previously identified direct target Plekha7 (Tavano et al., 2018) (Figure 4G-J and S4D-E), whose downregulation likely contributes to the transition from NSC to IPC. Finally, we observed a significant overlap of predicted targets with genes downregulated upon Insm1 overexpression in the E14.5 cortex (Tavano et al., 2018) (Figure 4K - 48/406, *p*<7.761e-17).

Collectively, these results highlight how integrating scRNA and scATAC and utilizing correlated enhancer-gene pairs can be used to successfully infer GRNs downstream of TFs, even in the absence of ChIP-seq validation or functional experiments. Using Eomes and Insm1 as examples, we could not only verify several known downstream genes, but also identify novel targets, significantly expanding our understanding of how TFs can modulate the chromatin landscape to regulate gene expression.

### Immuno-Methyl-Hi-C allows integration of single-cell epigenomics with bulk 3D genome organization and DNA methylation

To obtain a more comprehensive view of how chromatin landscape is reorganized during neural differentiation, we decided to employ a bulk approach using scRNA-based markers to sort precise cell populations, followed by combined mapping of 3D genome architecture and DNA methylation. Our method is based on the recently described methyl Hi-C protocols (Lee et al., 2019; Li et al., 2019), with the following improvements: 1) enrichment of desired cell types by immunoFACS; 2) significantly reduced input requirements (100-200k cells) and 3) higher proportion of Hi-C contacts/unique reads (Methods).

We decided to focus on the three major cell types identified using our scRNA-seq data and isolated NSC (Pax6^+^), IPC (Eomes^+^) and PN (Tubb3^+^) G_0_G_1_ cells from E14.5 somatosensory cortex in biological triplicates (Figure 5A, S5A-B). Our improved Methyl-Hi-C method was characterized by high bisulfite conversion efficiencies (Figure S5C-D), distance dependent decrease in contact probability (Figure S5E) and high reproducibility (Figure S5F). Furthermore, CpG methylation at gene bodies (Figure S5G) or Ctcf bound sites (Figure S5I) followed the expected pattern and did not change across cell types, indicating that global methylation levels remain the same.

**Figure 5.**
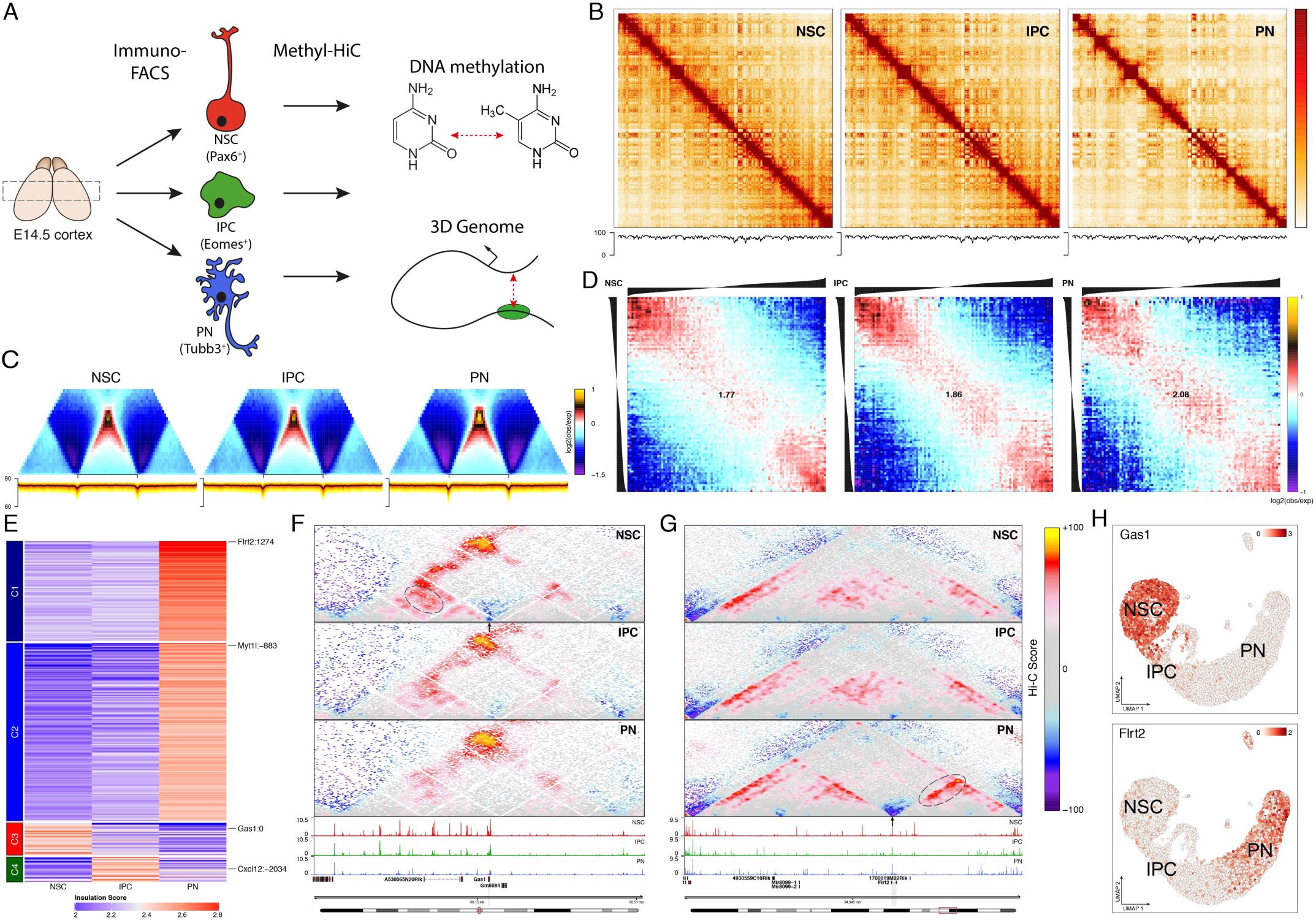
Immuno-methyl-Hi-C identifies global changes in 3D genome architecture during cortical development, independent of DNA methylation. (A) Schematic representation of the immunoFACS-based methyl-Hi-C experiment. (B) Knight-Ruiz balanced contact matrices for chr3 at 250 kb resolution (top) and DNA methylation levels (bottom). Scale bar is adjusted to account for the total coverage on chr3 in each cell type. (C) Average contact enrichment (top) and DNA methylation levels (bottom) across TADs. (D) Average contact enrichment between pairs of 250 kb loci arranged by their eigenvalue (shown on top). Numbers represent the compartment strength (Methods) (E) K-means clustering of differential TAD boundaries (n=322) based on the insulation score. (F) Contact maps (top) and aggregated accessibility of matched scATAC-seq clusters (bottom) for representative examples of a NSC- (F) or a PN- (G) specific TAD boundary (indicated by arrow) at the Gas1 or Flrt2 gene loci, respectively. Dynamic contacts are highlighted with dashed ellipse. (H) scRNA UMAP projection, colored by the expression levels of indicated genes.

Consistent with our previous results using an *in vitro* differentiation system (Bonev et al., 2017), we observed a global reorganization of chromatin interactions associated with fewer compartment transitions but stronger overall compartment strength (Figure 5B and 5D), as well as increased insulation at TAD boundaries and promoters upon neuronal differentiation (Figure 5C and S5H). However, this increase of insulation was not associated with changes in DNA methylation at TAD boundaries or accessibility/DNA methylation at CTCF bound sites (Figure 5C and S5H-I). We identified 322 differentially insulated domain boundaries (∼11% of all) (Figure 5E), many of which were in close proximity to TSS of cell-type-specific expressed genes such as Gas1 (NSC), Cxcd12 (IPC) or Flrt2 (PN) (Figure 5F-H, S5J), consistent with our previous findings (Bonev et al., 2017). Visual inspection of the identified loci highlighted several intra-TAD changes, which were frequently associated with dynamic enhancer accessibility (Figure 5F-G, S5J), prompting us to study the genome-wide dynamics of 3D connectivity and DNA methylation at the identified enhancer-gene pairs.

### Enhancer-gene pairs are associated with cell-type-specific changes in DNA methylation and chromatin looping

To address if there is a global rewiring of regulatory interactions, we first examined the aggregated Hi-C enhancer-promoter contacts, as previously described (Bonev et al., 2017; Methods). Strikingly, we found that: 1) positively correlated enhancer-gene pairs were characterized by increased contact strength compare to control (non-correlated) pairs and 2) interactions of positively correlated enhancer-gene pairs were much more cell-type-specific than control pairs with highest contact strength present in the cell type where the enhancer is the most active (Figure 6A and S6A). This effect was particular pronounced in NSC and PN, while IPC specific enhancer-gene pairs appeared to be in close proximity already in NSC. Nevertheless, IPC pairs increased in contact enrichment during the transition into IPC and decreased again in PN. These results suggest that although the majority of regulatory contacts can reform rapidly even between cells separated by only 1-2 cell divisions (NSC-IPC), there are some which are already pre-looped.

**Figure 6.**
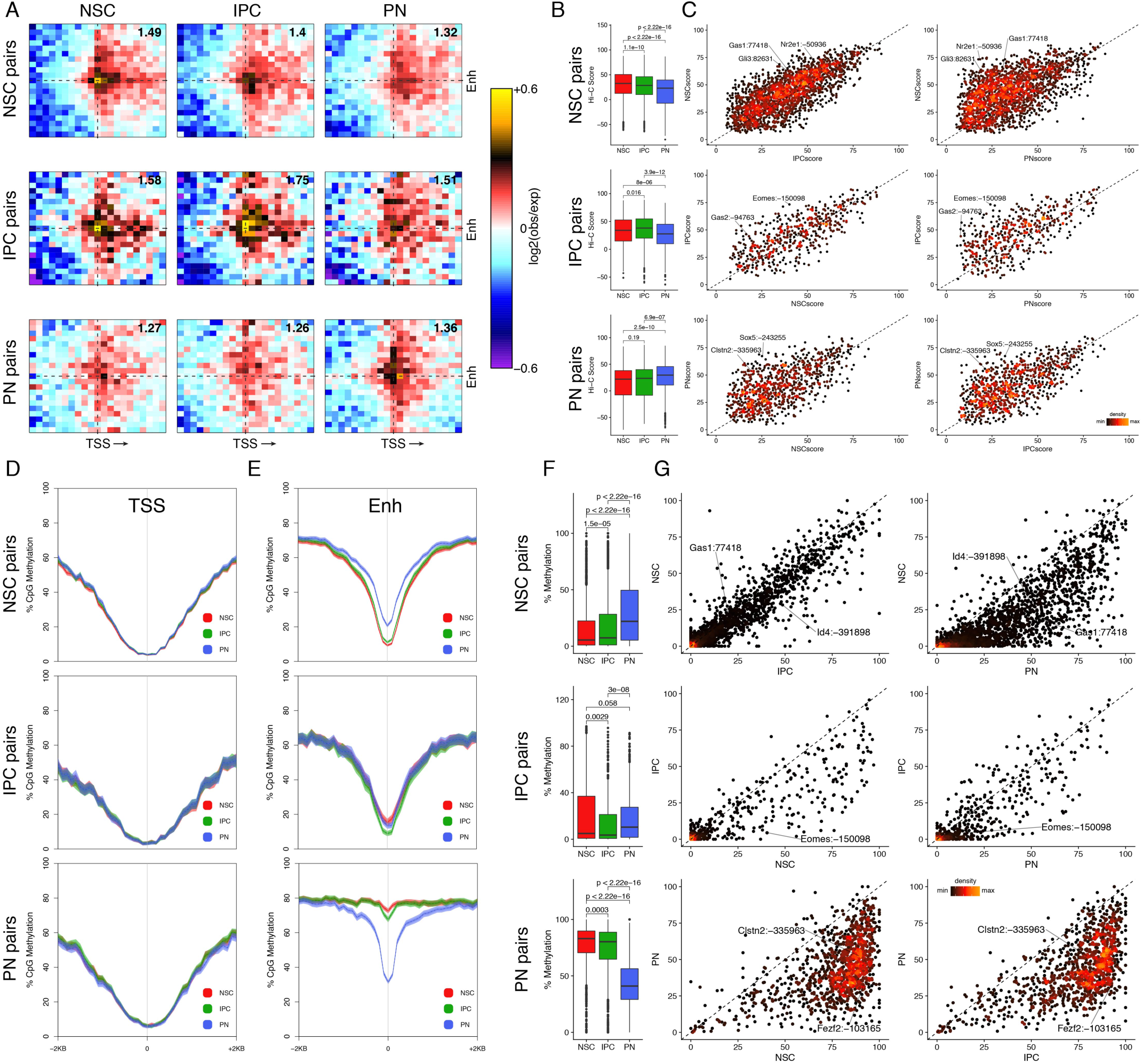
Chromatin loops and enhancer methylation levels are highly heterogeneous, yet mostly cell-type specific. (A) Aggregated Hi-C maps between enhancer (Enh) and the transcription start site (TSS) of NSC-, IPC- and PN-specific positively correlated enhancer-gene pairs. Genes are oriented according to transcription (arrow). Number in the top-right corner indicates the ratio of the center enrichment to the mean of the four corners (Methods). (B-C) Box-whisker (B) and scatter plots colored by density (C), depicting the Hi-C score at positively correlated cluster-specific enhancer-gene pairs. Statistical significance is calculated using wilcoxon rank-sum test. (D-E) Average DNA methylation levels at TSS (± 2kb) (D) and enhancer (± 2kb) (E) of positively correlated cluster-specific enhancer-gene pairs. Lines: mean values from biological replicates; semi-transparent ribbons: SEM (F-G) Box-whisker (F) and scatter plots colored by density (G), depicting the average DNA methylation levels at enhancers of positively correlated cluster-specific pairs. Statistical significance is calculated using wilcoxon rank-sum test.

To address this hypothesis directly, we examined the contact strength of each positively correlated or control enhancer-promoter pair independently for each cluster. First, we confirmed that only changes in contact strength in positively correlated, but not in control pairs were statistically significant (Figure 6B and S6B). Next, we plotted each pair separately and found, surprisingly, that individual enhancer-promoter pairs were characterized by a continuum of contact strengths ranging from strongly (Hi-C score ≥50) to weakly connected (Hi-C score ≤30) (Figure 6C). Examples of dynamic enhancer-promoter contacts include several cell-type-specific genes such as Gas1 (also depicted in Figure 5F), Gli3 and Nr2e1 in NSC; Eomes and Gas2 in IPC and neuronal-specific genes such as Sox5 and Clstn2. For some enhancer-promoter pairs the contact strength remained the same or even became stronger upon cell type transitions. Importantly, although non-correlated pairs were also characterized by high heterogeneity, they did not display the same bias in contact strength towards the cell type where the enhancer was active (Figure S6C).

To understand how DNA methylation is related to changes in enhancer accessibility and gene expression, we analysed the dynamics of CpG methylation at the identified enhancer-gene pairs. We found that promoter regions of the positively correlated linked pairs displayed consistent hypomethylation levels throughout neurogenesis (Figure 6D), in agreement with results in other differentiation systems (Meissner et al., 2008).

Contrary, DNA methylation levels at enhancers were lowest in cells where enhancers were most accessible (Figure 6E-F). This is fully consistent with previously described decrease of DNA methylation upon enhancer activation (Hahn et al., 2019; Noack et al., 2019; Stadler et al., 2011; Zhu et al., 2016). Although the majority of enhancers became hypomethylated upon activation, there was considerable heterogeneity in DNA methylation levels (Figure 6G). Non-correlated enhancers were associated with higher overall levels of DNA methylation and were not as dynamic (Figure S6D-F). Interestingly, although chromatin interactions between anti-correlated pairs did not change significantly, DNA methylation levels varied between cell types, suggesting that these two molecular layers are not strictly correlated (Figure S6G-H).

In summary, we showed that, overall, regulatory interactions between correlated enhancer-gene pairs are fairly dynamic and strongest in the cell type where the enhancer is active and the linked gene expressed. Nevertheless, there is a considerable heterogeneity at the level of individual enhancer-promoter pairs: while many follow the classical model of cell-type-specific looping, some appear to be already pre-looped before transcription occurs and others are only weakly interacting with their target promoters. DNA methylation levels at enhancers are generally anti-correlated with accessibility but, again, vary considerably from enhancer to enhancer. Importantly, both of these metrics, 3D contact strength and DNA methylation, could not be simply inferred from scRNA/scATAC, highlighting the importance of using multiple epigenome layers to study GRNs.

### Epigenome remodelling is associated with tissue-specific transcription factors

Given the observed dynamics and heterogeneity at the identified enhancer-gene pairs in both DNA methylation and interaction strength, we sought to identify the potential molecular mechanisms underlying these observations. Transcription factors have been previously suggested as potential mediator of such processes by us and others (Bonev et al., 2017; Hahn et al., 2019), but it is unclear which and to what extend they participate in these biological processes.

To address this question, we developed a computational method to assess the average interaction strength and cellular specificity for each TF based on accessible sites containing their binding motif (Methods). We first verified the method using pairs of convergent Ctcf sites as control (Rao et al., 2014), which had the highest maximum normalized Hi-C score as expected (Figure 7A – Ctcf_ForRev). Surprisingly, we found several key neurogenic factors which have not been previously described in the context of chromatin loop formation (Figure 7A and S7A). These include the basic helix-loop-helix TF Neurog2, as well as members of the POU-domain-class3 (Pou3f2, Pou3f3) and Sox (Sox2, Sox4, Sox8) families. Interestingly, in addition to high interaction strength, these factors were also characterized by the highest variance, suggesting that strong interactions were formed transiently in a cell-type-specific manner. To confirm these findings, we used two different approaches. First, we plotted the aggregated Hi-C contact maps for Pou3f2 motif containing peaks and observed high interaction in NSC and IPC followed by a drop in PN where Pou3f2 is downregulated (Figure 7B-C). Similarly, contact strength of Neurog2 motif containing peaks was the highest in IPC, which correlated well with its expression pattern (Figure 7B-C). Second, we used ChIP-seq based peaks to confirm the cell-type-specificity between Neurog2/Eomes bound sites, as well as Pax6, as previously described (Figure S7B; Bonev et al., 2017). This dynamic pattern of chromatin interaction matched well with the changes in accessibility at TF bound sites (Figure S7C).

**Figure 7.**
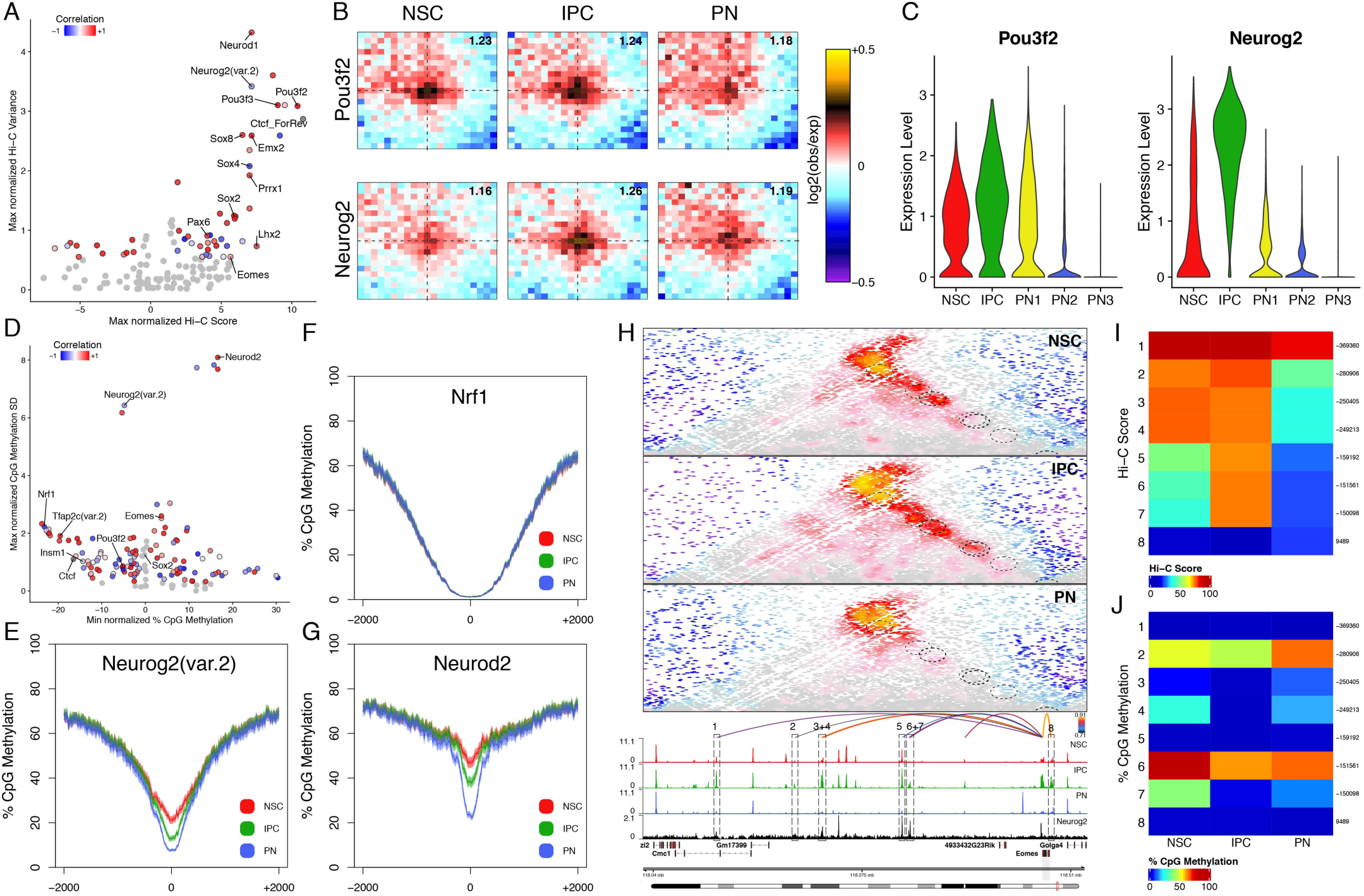
Neurog2 is associated with changes in both chromatin looping and DNA methylation levels. (A) Scatterplots depicting the maximum Hi-C score and the normalized variance at pairs of accessible peaks associated with specific TF binding motifs. TFs are colored based on the Pearson correlation between their expression and the accessibility of their binding motif. Grey circles represent non-significant TFs (p>0.05; permutation test). Only TFs where correlation between expression and motif accessibility can be calculated (Figure 3E) are considered. (B) Aggregated Hi-C plots between intraTAD pairs of accessible peaks overlapping with the indicated TF motif. Number in the top-right corner indicates the ratio of the center enrichment to the mean of the four corners (Methods). (C) Violin plots depicting the expression levels of indicated TFs per scRNA cluster. (D) as in (A) but comparing the minimum DNA methylation level and the normalized SD at accessible peaks associated with specific TF binding motifs (E-G) Average DNA methylation levels at accessible peaks overlapping with the indicated TF binding motifs. Lines: mean values from biological replicates; semi-transparent ribbons: SEM (H) Contact maps (top) and aggregated accessibility of matched scATAC-seq clusters (bottom) at the Eomes locus. Depicted are the identified linked enhancers (1-8, arcs) and Neurog2 ChIP-seq track (Sessa et al., 2017). Arcs on top are colored by Pearson correlation of the enhancer accessibility and Eomes expression. (I-J) Heatmaps depicting the Hi-C score of linked enhancer-Eomes pairs (I) or the DNA methylation levels at the corresponding Eomes enhancers (J).

Next, we asked if we could use an analogous approach to identify TFs related to dynamic DNA methylation (Methods). Among those predicted to be associated with low levels of DNA methylation, we found Nrf1 (Figure 7D-F), which has been previously characterized to be highly sensitive to DNA methylation levels (Domcke et al., 2015), thus validating our approach. Neurod2 and Neurog2 were some of the TFs with the highest variance (Figure 7D-E and G) and while Neurod2 has been previously shown to be important for establishing cell-type-specific DNA methylation in the cortex (Hahn et al., 2019), not much is known about the role of Neurog2 in this process. We confirmed these findings using positively correlated enhancer-gene pairs as well as Neurog2/Neurod2 peaks (based on ChIP-seq data), instead of motifs (Figure S7D-E).

Finally, we focused on our attention on Neurog2, as it was one of the few TFs which showed high cell specificity in both chromatin interactions and DNA methylation, and due to its well-characterized role in neuronal differentiation in the cortex (Mattar et al., 2008; Schuurmans et al., 2004). Using the linkageScore method described earlier, we identified Eomes as a putative target of Neurog2 (Figure S7F). Visual inspection at the Eomes locus showed a complex regulatory network with at least 8 positively correlated enhancer-gene pairs (Figure 7H). Importantly, enhancers bound strongly by Neurog2 engaged in strong, cell-type-specific looping with the Eomes promoter only in IPCs (Figure 7H-I, loci 5-7), while other enhancers were either pre-looped (Figure 7H-I, loci 1-4) or not interacting strongly (Figure 7H-I, locus 8). DNA methylation levels at Neurog2 bound enhancers were either very low (Figure 7J, locus 5) or decreased specifically in IPC (Figure 7J, loci 4, 6 and 7).

Collectively, these experiments identify the previously underappreciated role of transcription factors in dynamic chromatin looping and DNA methylation. Furthermore, several novel TFs with a well-characterized role in cortical development such as Neurog2, Pou3f2 and Eomes are associated with strong cell-type-specific looping, but only Neurog2 binding is correlated with changes in DNA methylation. Finally, we identify Eomes as a potential downstream target of Neurog2 and highlight the complex regulatory network of pre-looped and dynamic enhancers involved in regulating its expression.

## Discussion

The identification of lineage specific gene regulatory networks during development has remained challenging, in particular in complex, heterogeneous tissues *in vivo.* Another layer of complexity derives from the difficulty to simultaneously profile the multiple level of molecular regulation at play. Here, we combined single-cell transcriptomics and epigenomics with cell-type-specific bulk DNA methylation and 3D genome architecture *in vivo* to address how the chromatin landscape is reorganized during cortical development. Our rich datasets and the novel computational methods for data integration provided numerous insights into crucial cell fate transitions during neural differentiation in the embryonic cortex.

We observed that during neuronal differentiation cells transition along a continuum of cellular states, in contrast with the defined cell types present in the mature postnatal cortex (Zeisel et al., 2015). Using RNA velocity and trajectory analysis approaches, we confirmed the dynamic expression of several known TFs (Telley et al., 2016) and identified novel temporally-regulated genes related to chromatin remodeling (Chd7, Rbbp7) or associated with autism and mental retardation (Lrfn5). Our findings confirm and significantly advance initial single-cell data (Loo et al., 2019; Telley et al., 2016, 2019), which were based on a small cohort of cells and/or low sequencing depth.

Our matched scATAC-seq data represents one of the highest-quality scATAC datasets currently available (46899 unique fragments/cell), paving the way to comprehensively characterize the dynamics of the chromatin landscape in neuronal differentiation. As suggested by other model systems (Corces et al., 2016; Klemm et al., 2019), promoter accessibility is generally not correlated with gene expression, while distal elements and transcription factor motifs are characterized by high variability and specificity.

Using an integration approach pioneered by the Greenleaf (Granja et al., 2019, 2020) and Satija (Stuart et al., 2019) labs, we identified thousands of positively correlated enhancer-gene pairs, generating a rich resource to study GRNs in the cortex. We found a previously unexpected connection between expression of cell-type-specific TFs and the accessibility at the enhancers overlapping their binding motif, indicating that many TFs potentially function by remodeling the chromatin landscape. Whether or not binding of these TFs is a cause (thus challenging the concept of only a limited number of pioneer TFs) or a consequence of changes in enhancer accessibility, remains to be seen. Using an integrated trajectory analysis, we showed that enhancer activation generally precedes transcription at dynamic genes, although only a subset of enhancers appears to be truly lineage-priming. Our findings corroborate prior conclusions in early mammalian embryogenesis (Argelaguet et al., 2020) and recent data using simultaneous paired ATAC/RNA measurements in single cells (Ma et al., 2020).

3D genome architecture and DNA methylation represent two additional less characterized molecular layers of the regulatory landscape, but have only rarely been coupled to single-cell epigenomic measurements (Clark et al., 2018). Our immuno-FACS approach allowed us to combine the resolution and depth of bulk Hi-C/DNA methylation with cell-type-specificity. The reduced input requirements and the improved resolution compared to prior studies (Lee et al., 2019; Li et al., 2019) enable the application of this method to other tissues and model organisms, where high-quality antibodies exist.

Previous studies have produced somewhat conflicting results regarding the importance of 3D genome architecture on gene regulation(Deng et al., 2012; Ghavi-Helm et al., 2019; Llinares-Benadero and Borrell, 2019; Lupiáñez et al., 2015; Nora et al., 2017) and the cell-type-specificity of regulatory loops (Bonev et al., 2017; Ghavi-Helm et al., 2014; Javierre et al., 2016). Our results indicate that contact strength between enhancers and promoters (as well as DNA methylation levels) is not simply a consequence of gain in accessibility/expression and cannot be inferred from other type of measurements (such as matched accessibility/gene expression), as even highly correlated enhancer-gene pairs were characterized by a continuum of interactions scores. Despite this apparent heterogeneity, there are clear genome-wide trends in the data, suggesting that the majority of enhancer-promoter interactions are dynamic and cell-type-specific, although such differences are rather quantitative than binary (loop vs no loop).

Finally, we provide evidence that a subset of transcription factors is associated with strong and cell-type-specific chromatin looping, potentially acting as “molecular bridges” to facilitate the reorganization of the epigenetic landscape in development. We identify the proneural TF Neurog2 as one of the few factors correlating with both dynamic 3D contacts and DNA methylation changes and characterize the chromatin landscape at its predicted downstream targets Eomes and Rnd2.

Collectively, our data provides a comprehensive view of the reorganization of the epigenetic regulatory landscape during lineage commitment in the mouse cortex. The novel insights gained from our integrative analysis can be used to more precisely define, compare and ultimately engineer cellular identities both in development and evolution, as well as for therapeutic and regenerative purposes. Our results pave the way for functional studies aiming to resolve the relative influence of dynamic vs pre-looped enhancers on gene expression and the importance of transcription factor binding to the rewiring of the regulatory 3D genome. Finally, our data provides a rich resource to study the reorganization of chromatin, the transcriptomic landscape and the establishment of lineage-specific GRNs in cortical development.

## Author Contributions and Notes

F.N. and B.B. conceptualized the study. F.N. performed the single-cell and the methyl-Hi-C experiments and contributed to the data analysis. S.V. performed the immunoFACS experiments. M.C. performed the RNAscope and IF experiments. F.C contributed to the single-cell data analysis. B.B assisted with experiments, developed the computational framework to analyze the single-cell and methyl-Hi-C data and supervised the project. F.N and B.B wrote the manuscript with input from all authors. The authors declare no competing interests.

## Acknowledgments

We thank Prof Magdalena Götz, Prof Giacomo Cavalli and Dr Thomas Schwarz-Romond for critically reading the manuscript. We thank Jei Diwakar for assisting with the graphical abstract and the members of the Bonev and Götz lab for the useful discussions. Sequencing was performed at the Helmholtz Zentrum München (HMGU) by the NGS-Core Facility and cell sorting was done at the Flow Cytometry core facility at the Biomedical Center, LMU. Work in B.B. group was supported by the Helmholtz Pioneer Campus, DFG priority program SPP2202 (BO 5516/1-1) and the Helmholtz-Gemeinschaft (ERC-RA-0037).

## Materials and Methods

### Experimental Model and Subject Details

Time-mated pregnant C57BL/6JRj mice were obtained from Janvier Laboratories (Route du Genest, 53940 Le Genest-Saint-Isle, France) and kept under standard housing conditions according to local regulations of the Regierung Oberbayern, Germany. E14.5 mouse embryos were used sex independent. Each biological replicate represents a single embryo from different mothers in case of scRNA-seq/scATAC-seq or a pool of 4-6 littermates from separate mothers in case of MethylHiC. All experiments were performed according to national guidelines and where approved by local authorities (Regierung Oberbayern, Germany: ROB-55.2-2532.Vet_02-19-175).

### Experimental procedure

#### Tissue preparation and dissociation

Brains of E14.5 embryos were either used directly for in-situ hybridization and immunohistochemistry or were further dissected to isolate the somatosensory cortex with a prior removal of all meninges. The dissected cortex was dissociated using a papain-based neural dissociation kit (Miltenyi Biotec, Cat. N: 130-092-628) according to the manufacturer protocol with minor modifications. Briefly, samples were spun down with 300g for 2min. at RT, supernatant was removed and samples were mixed and incubated with prewarmed enzyme mix 1 for 15min. at 37°C under slow rotation. Enzyme mix 2 was added and gently mixed using a disposable Pasteur pipette (ThermoFisher, Cat. N: PP89SA). Subsequently, samples were incubated two times for 5min. at 37°C under slow rotation with a manual dissociation in between using a disposable Pasteur pipette. The single cell suspension was passed twice through a 40µM cell strainer (VWR, Cat. N: 734-2760), spun down with 300g for 5min. at 4°C and washed twice with ice cold HBSS (ThermoFisher, Cat. N: 14025092). After the final wash cells were resuspended in PBS with 1% BSA (ThermoFisher, Cat. N: AM2618) and the cell number as well as viability were assessed using Countess™ II Automated Cell Counter (Invitrogen).

#### scRNA-seq and scATAC-seq

Directly after tissue dissociation scRNA-seq (v3, 10x Genomics, Cat. N: PN-1000075, PN-1000074, PN-120262) as well as scATAC-seq (10x Genomics, Cat. N: PN-1000110, PN-1000086, PN-1000084) libraries were generated according to the manual instructions with a targeted recovery of 6000 cells/nuclei per sample.

#### ImmunoFACS

Dissociated cells were fixed for 10min. at RT in 1% freshly prepared Formaldehyde in PBS (ThermoFisher, Cat. N: 28908) under slow rotation and quenched by addition of Glycine (Invitrogen, Cat. N: 15527013) to a final concentration of 0.2M. Fixed cells were spun down with 500g for 5min. at 4°C, washed once with 1% BSA, 0.1% RNAsin plus RNase inhibitor (Promega, Cat. N: N261A) in PBS (wash buffer) and subsequently incubated for 10min. at 4°C in permeabilization buffer consistent of 0.1% freshly prepared Saponin (Sigma-Aldrich, Cat. N: SAE0073), 0.2% BSA (ThermoFisher, Cat. N: 15260-037) and 0.1% RNAsin plus RNase inhibitor in PBS. The permeabilization buffer was removed by centrifuging the cells with 2500g for 5min. at 4°C followed by staining against Pax6 (1:40; BD Bioscience, Cat. N: 561664), Eomes (1:33; BD Bioscience, Cat. N: 566749) and Tubb3 (1:14; BD Bioscience, Cat. N: 560394) in staining buffer (0.1% saponin, 1% BSA, 0.1% RNAsin plus RNase inhibitor in PBS) for 1h at 4°C under slow rotation. Cells were washed twice with permeabilization buffer, once with wash buffer containing DAPI (1:1000; ThermoFisher, Cat. N: 62248) and a final wash with wash buffer without DAPI. Between each washing step, the cells were incubated for 5min. at 4°C with the respective buffer under slow rotation and spun down with 2500g for 5min. at 4°C. After the last wash the cells were resuspended in PBS with 1% BSA and 1% RNAsin plus RNase inhibitor, passed through a 40µM cell strainer (ThermoFisher, Cat. N: 15342931) and immediately FAC-sorted.

#### Fluorescence-activated cell sorting

FAC-sorting was performed on a BD FACSAria Fusion (BD Bioscience) with four lasers (405, 488, 561, 640) using a 100µm nozzle. After selecting singlets using forward and side scatter, cells in G_0_G_1_ were identified by genomic content based on DAPI staining. Subsequently, these cells were divided into Tubb3 high for PN and low for progenitor cell types. The progenitor population was further subdivided into Pax6-high/Eomes-low for NSC and Eomes-high for IPC. The set gates are displayed in Figure S5A. After sorting, cells were either directly pelleted (500g for 5min. at 4°C), flash frozen in liquid nitrogen and stored at -80 °C for further usage or RNA was extracted using the Quick-RNA FFPE Miniprep kit (Zymo Research, Cat. N: R1008) with Zymo-Spin IC Columns (Zymo Research, Cat. N: C1004-250).

#### Real Time Quantitative PCR (qPCR)

Reverse transcription was performed using Maxima H Minus Reverse Transcriptase (ThermoFisher, Cat. N: EP0751) with OligodT primer (ThermoFisher, Cat. N: SO132) according to the manual instructions. Transcripts were quantified by using LightCycler® 480 SYBR Green I Master Mix (Roche, Cat. N: 04707516001) with the appropriate primers (see key resource table) on a Roche LightCycler® 480.

#### Methyl-Hi-C

For Methyl-Hi-C we adapted current protocols (Lee et al., 2019; Li et al., 2019). Briefly, frozen pellets of fixed cells were thawed on ice, lysed with 0.2% Igepal-CA630 (Sigma-Aldrich, Cat. N: I3021), permeabilized with 0.5% SDS (Invitrogen, Cat. N:AM9823) and digested with 200U DpnII (New England Biolabs, Cat. N: R0543) overnight at 37°C. Subsequently, sticky ends were filled by incubating the nuclei for 4h at RT with DNA Polymerase I (New England Biolabs, Cat. N: M0210) and a biotin-14-dATP (Life Technologies, Cat. N: 195245016) containing nucleotide mix in DpnII buffer followed by proximity ligation for at least 6h at 16°C. Thereafter nuclei were lysed, proximity ligated DNA was reverse-crosslinked overnight at 68°C followed by a purification by ethanol precipitation. DNA was sheared to ∼550 bp fragments using a Covaris S220 sonicator and end-repaired by incubating the samples with T4 DNA Polymerase (New England Biolabs, Cat. N: M0203) for 4h at 20°C. Prior to bisulfite conversion, sheared and biotinylated fully methylated pUC19 (Zymo Research, Cat. N: D5017) and unmethylated lambda DNA (Promega, Cat. N: D1521) was added to the samples (∼ 0.01%). Bisulfite conversion was performed using the EZ DNA Methylation-Gold kit (Zymo Research, Cat. N: D5005) followed by library construction using the Accel-NGS® Methyl-Seq DNA Library kit (Swift Bioscience, Cat. N: 30024) according to the manufacturer’s instructions until the adapter ligation step. After this step, biotin pulldown was performed using MyOne Streptavidin T1 beads (ThermoFisher, Cat. N: 65602) followed by 5 washes with washing buffer containing 0.05% Tween-20 (Sigma-Aldrich, Cat. N: P9416) and two additional washes with low-TE water. Libraries were amplified on the streptavidin beads using the EpiMark Hot Start Taq (New England Biolabs, Cat. N: M0490) using following program: 95°C 30s; {95°C 15s, 61°C 30s, 68°C 60s} x14; 68°C 5min; Hold at 10°C. Streptavidin T1 beads were pelleted on a magnetic rack and the prepared libraries within the supernatant were purified using 0.6x AMPure XP beads (Agencourt, Cat. N: A63881) to reach an average fragment size of approximately 500bp.

#### Library QC and Sequencing

Before sequencing, libraries were quantified by qPCR using either the NEBNext® Library Quant kit (New England Biolabs, Cat. N: E7630S) or the KAPA Library Quantification Kit (scRNA-seq libraries only; Roche, Cat. N: 07960298001). Size distribution of the obtained libraries was assessed using Agilent 2100 Bioanalyzer. Libraries were sequenced 2×100bp paired end on an Illumina NovaSeq 6000 to depth of approximately 400 million PE reads per replicate for scRNA-seq, 473 million PE reads per replicate for scATAC-seq and 320 million PE reads per replicate for MethylHiC.

#### Sectioning, Immunohistology and In-situ hybridization

Brains were fixed in freshly prepared 4% Formaldehyde in PBS at 4°C for at least 8h, washed once in PBS and cryoprotected in 30% sucrose for ∼6h at 4°C. Subsequently, brains were embedded in TFM-Tissue Freezing Media (TBS-Triangle Biomedical, Cat. N: 15-183-13), snap-frozen on dry ice and finally cryosectioned (∼10μm) using a CryoStar NX70 (ThermoFisher). Sections were collected on Superfrost Plus adhesive microscope slides (ThermoFisher, Cat.N: 7608105) and stored at −80°C until further use.

For immunohistochemistry, the sections were hydrated in PBS and incubated for 30min. at RT in 0.1M Glycine (Invitrogen, Cat.N: 15527013) in PBS followed by two washes with PBS. Subsequently sections were incubated for 1h at RT in PBS blocking buffer containing 5% horse serum (Sigma Aldrich, Cat. N: H0146), 0.3% Triton X-100 (Sigma Aldrich, Cat. N: X100). Staining was performed overnight at 4°C with anti-Pax6-A488 (1:200, BD Biosciences, Cat. N: 561664), anti-Eomes-PE (1:200, BD Biosciences, Cat. N: 566749) and anti-Tubb3-A647 (1:200, BD Biosciences, Cat. N: 560394) antibodies in PBS blocking buffer. Section were washed three times for 10min with 0.1% Triton X-100 in PBS, followed by an incubation with DAPI (1:1000 diluted in PBS) for 10min. and finally mounted using Fluoromount-G (Invitrogen, Cat. N: 00-4958-02).

RNA In-situ hybridization was performed using the RNAscope Multiplex Fluorescent Reagent kit v2 (ADCBio, Cat. N: 323100) according to the manual instructions. Briefly, slides were baked 30min. at 60°C, dehydrated in ethanol and pre-treated with hydrogen peroxide for 10min. at RT. Antigen retrieval was performed for 5min. at ca. 98°C, followed by protease incubation 15min. at 40°C. Probe hybridization was done for 2h at 40°C. Signal amplification and detection reagents, including Opal fluorophores (Akoya Biosciences, Cat. N: FP1487001KT, FP1488001KT, FP1497001KT), were applied sequentially. Nuclei were counterstained with DAPI and slides mounted with Fluoromount-G (Invitrogen, Cat. N: 00-4958-02).

All images were acquired using a Zeiss LSM 710 confocal microscope.

### scRNA-seq

#### scRNA-seq processing

Raw sequencing data was converted to fastqs using cellranger mkfastq (10x Genomics, v3.1.0). scRNA reads were aligned to the GRCm38 reference genome (mm10) and quantified using cellranger count (10x Genomics v3.1.0).

#### scRNA-seq quality control

We removed low-informative cells by filtering cells with less than 1000 genes or 2500 UMIs per cell detected. To lower doublet representation we filtered cells with more than 7000 genes per cell detected and the top 4% cells (estimated doublet percentage) with the highest number of UMIs. Finally, to remove any potential dead cells we filtered cells that have more than 10% mitochondrial counts.

#### scRNA-seq clustering and dimensionality reduction

Seurat v3.1.5 (Stuart et al., 2019) was used to further process the cells passing the QC filters. After log transformation, feature selection using variance stabilizing transformation (top 2000 most highly variable genes) and linear transformation, principal component analysis (PCA) was performed using the first 20 dimensions. After dimensionality reduction, Harmony (Korsunsky et al., 2019) was used to correct the batch effect between the two biological replicates. Next, we applied Louvain clustering with resolution 0.3, n.start=100 and n.iter=500 and visualized the data using UMAP (min.dist=0.5, spread=1, n.epochs=500). As they represent very few cells, the microglia and mural clusters were further manually identified based on the UMAP projection and the subclustering of the NSC cluster. Cluster identify was determined based on the top40 differentially expressed genes (MAST – (Finak et al., 2015), min. log fold change of 0.25 and expressed in at least 25% of the cells in the cluster)

#### scRNA-seq velocity and pseudotime analysis

The percentage of spliced and unspliced reads was calculated using Velocyto v0.17 (La Manno et al., 2018) and RNA velocity was calculated using scVelo (dynamical model) (Bergen et al., 2020). Only cells passing the previously described QC were used and UMAP coordinates were transferred from Seurat. To calculate trajectory and pseudotime, we used Monocle3 (Cao et al., 2019), while retaining cluster assignment and UMAP coordinates from Seurat. Trajectory graph was constructed using the following parameters: minimal_branch_len=20, ncenter=600, geodesic_distance_ratio=0.275 and cells belonging to the NSC cluster were assigned as root cells. To calculate the change of gene expression as a function of the pseudotime, we fitted a generalized additive model using cubic regression splines and REML smoothing for each of the top 3000 most variable genes (expressed in at least 20 cells). The values were then rescaled per gene from 0 to 1.

### scATAC-seq

#### scATAC-seq processing

Raw sequencing data was converted to fastqs using cellranger-atac mkfastq (10x Genomics, v1.2.0). Reads were aligned to the GRCm38 reference genome (mm10) and quantified using cellranger-atac count (10x Genomics v1.2.0) using integrated doublet removal.

#### scATAC-seq quality control

We calculated the QC statistics separately per replicate and filtered the combined 10x object (merged using cellranger-atac-aggr with no normalization). To ensure sufficient sequencing depth and high signal-to-noise ratio, we filtered cells with less than 10000 unique fragments per cell and TSS enrichment ratio less than 8 or more than 25 (Figure S2B). To account for any remaining doublets after the automatic cellranger-atac filtering, we additionally removed cells with more than 120000 unique fragments per cell. TSS enrichment was calculated as described (Granja et al., 2019). In brief, Tn5 insertions located within ±2000 bp relative from each TSS (strand-corrected) were aggregated per TSS, normalized to the mean accessibility ±1900-2000 bp from the TSS and smoothed every 51bp. Maximum smoothed value was reported as TSS enrichment. Fragment size distribution for cells passing the QC filters was calculated using ArchR (Granja et al., 2020) and plotted with ggplot2.

#### scATAC-seq clustering and dimensionality reduction

To obtain a set of initial clusters we first counted the number of unique fragments in 5 kb genomic bins (Signac– https://github.com/timoast/signac). After binarization, the top20000 accessible windows were kept and the matrix was transformed using log term frequency-inverse document frequency (TF-IDF) transformation using Signac. The normalized matrix was then used as an input for partial singular value decomposition (SVD) using irlba (Signac). After dimensionality reduction, Harmony (Korsunsky et al., 2019) was used to correct the batch effect between the two biological replicates. Next, we retained the first 20 dimensions, applied Louvain clustering with resolution 1, n.start=50 and n.iter=50 and visualized the data using UMAP (min.dist=0.5, spread=1.5, n.epochs=2000). For each cluster, peak calling was performed on the Tn5-corrected insertions as described in Granja et al. 2019. Peak size was then normalized to 501 bp length, filtered by the mm10 ENCODE blacklist and then peaks were merged into a union set as previously described (Granja et al., 2019). Next, this high-quality peak set was used to generate the final clustering and visualization. First fragments containing within peaks were calculated using Signac, binarized and the top 25000 variable peaks were identified (using aggregated counts per million from the initial bin-based clusters). The count matrix associated with those peaks was then again subjected to TF-IDF normalization followed by SVD as described above. After batch correction using Harmony (Korsunsky et al., 2019), the first 20 dimensions were retained and clusters were identified using Louvain algorithm (resolution=0.3, n.start=100, n.iter =200). Data was visualized using UMAP embedding (min.dist=0.5, spread=1.5, n.epochs=2000). Cluster identify was determined based on the top40 differentially accessible gene bodies (Student’s t-test, min log fold change of 0.25 and expressed in at least 25% of the cells in the cluster).

#### scATAC-seq calculation of promoter and gene body

To calculate gene body accessibility scores, we counted the number of unique fragments along the whole span (TSS – TTS) of protein coding genes (EnsDb.Mmusculus.v79), extended 2000 upstream of TSS. To calculated promoter scores, we counted the number of unique fragments along promoters of protein coding genes (defined as the sequence −2000 bp to +200 bp of the TSS.

#### scATAC-seq motif and ChIP-seq accessibility deviations

Motif accessibility was calculated using chromvar (Schep et al., 2017) as implemented in the Signac package (https://github.com/timoast/signac). In brief, position-weight-matrices were obtained from the JASPAR2020 motif database (Fornes et al., 2020), to which entries present in the JASPAR2018 version but subsequently removed, were manually added. Each accessibility peak was then tested for the presence/absence of each transcription factor motif and GC-bias-corrected deviations were computed using the chromVAR ‘deviations’ function as implemented in Signac (‘RunChromVar’). Accessibility deviations associated with ChIP-seq peaks were computed analogously, but the overlap of ChIP-seq peaks and scATAC peaks was used as entry to chromVAR instead.

#### scATAC-seq unique peaks identification

Cluster-specific peaks were identified using feature binarization as described (Granja et al., 2019). In brief, pseudobulk replicates were created for clusters with N cells < 100, while real biological replicates were used for the remaining clusters. Peaks were considered as unique if they had adjusted P value less than 0.01 and minimum log fold-change of 0.25 to the next highest cluster. The identified unique peaks were split into two categories: promoter-associated (less than 500 bp away from a TSS) and distal (more than ±5 kb away from an annotated TSS).

#### TF footprinting

The Tn5-normalized accessibility around TF motifs was calculated as previously described (Corces et al., 2018) using the ArchR package (Granja et al., 2020). The expected Tn5 bias was substracted from the calculated footprints to generate the final footprint plots.

### scRNA and scATAC-seq integration

#### Label transfer and co-embedding

To integrate scRNA and scATAC datasets we used Seurat’s CCA (Stuart et al., 2019). In brief, first we identified transfer anchors using the top 5000 most variable genes shared across both datasets using FindTransferAnchors (dims=1:20, k.anchor=20, k.filter=200, k.score=30, max.features=500). We then transferred the scRNA based labels using the inferred anchors and the harmony corrected low-dimensional coordinates as weight reduction. After we confirmed the high-confidence of the prediction scores (Figure S3B), we then co-embedded the scRNA and the scATAC cells in the same low-dimensional space and recalculated UMAP embedding (Figure S3A, dims=1:20, n.epochs=2000, spread=1.5, min.dist=0.5). To enable more robust downstream correlation-based analysis we used the previously described Cicero-based kNN approach to group scATAC-seq accessibility (4892 groups, KNN=50) match grouped gene expression (based on scRNA-seq closest neighbors).

#### Identifying pairs of matched genes and predicted enhancers

To identify putative enhancers, where the accessibility of the predicted distal regions correlated with changes in gene expression (and not accessibility of the promoter), we adapted the approach first described by Granja et al. 2019. Briefly, the correlation between the logNormalized matched grouped scATAC and scRNA values was calculated for each pair of distal scATAC peak (at least 5kb away from any annotated TSS) and gene promoter within a maximum genomic distance of 500 kb. The significance of the calculated correlations was determined using a trans-based null correlation and peak-to-gene links with significance of FDR<0.1 and Pearson correlation of > 0.35 were considered as positively correlated (referred to as putative enhancer-promoter pairs for simplicity in the text). In addition, we also identified two additional classes of peak-to-gene relationship: negatively correlated (FDR<0.1 and r< −0.35) and control pairs (FDR>0.1 and −0.35 < r < 0.35). As the number of control pairs is much higher than either positively or negatively correlated once, we subsampled this category to match the positively correlated pairs. Cluster-specific pairs were determined using pseudobulk feature binarization as described above.

#### Integrated pseudotime analysis

Pseudotime on the combined integrated scRNA-scATAC object was calculated using Monocle3, analogous to scRNA-seq alone. Imputed gene expression values based on gene body accessibility and integration vectors as described previously were used together with measured scRNA values to construct a cellDataSet object, retaining cluster assignment (scRNA-based) and UMAP coordinates from Seurat. To calculate the change of accessibility and motifs deviations as a function of the pseudotime, we fitted a generalized additive model using cubic regression splines and REML smoothing, analogous to gene expression for scRNA-seq. The values were then rescaled per gene from 0 to 1. For each enhancer-gene pair we then ordered the cells based on their pseudotime and calculated the pseudotime difference between their respective maxima, which was used to infer the relationship between enhancer accessibility and gene expression (Figure 3D). Analogous process was repeated for TF motif deviations and the correlation between the gene expression pattern of each TF and its corresponding motif accessibility along the pseudotime was computed as well.

#### Identification of predicted TF targets

We reasoned that we can predicted direct targets of TF in the absence of available ChIP-seq data based on the enrichment of the TF motif in the positively correlated enhancer-gene pairs. First, we identified all enhancer-gene pairs which contained the corresponding TF motif (either in the distal region or the promoter region). We then calculated gene “linkageScore” by adding up the r^2^ from each pair per gene (if the motif was contained in the promoter, we used a value of r=1). To calculate motif enrichment, we used background peaks with similar GC content and determined significance using hypergeometric test. Although this approach works well for genes with multiple links, the significance of motif enrichment for genes with very few identified pairs cannot be accurately calculated.

### Methyl-Hi-C analysis

#### Mapping and QC

We mapped the joint Hi-C / DNA methylation data to the mm10 genome using JuiceMe (Durand et al., 2016). Only uniquely mapping reads (mapq>30) were retained for further analysis. After removal of PCR duplicates, reads were translated into a pair of fragment-ends (fends) by associating each read with its downstream fend. CpG methylation was assessed using MethylDackel (https://github.com/dpryan79/MethylDackel), in a “mergeContext” mode with the first 6 nucleotides omitted from further analysis. Reads from individual replicates were pooled and only Cs in a CpG context with at least 10x total coverage were further analysed. For Hi-C, reads mapping to the same restriction fragment or separated by less than 1 kb were excluded from further analysis. The QC metrics were reported in Supplementary Table S1.

#### QC of bisulfite conversion efficiency

To determine the efficiency of the bisulfite conversion we determined the proportion of CpG methylation which was detected in fragments mapping to the lambda DNA sequence. On average this represents ∼0.5%, suggesting 99.5% conversion rates (Figure S5C). To calculate the detection rate we determined the proportion of CpG on fully methylated pUC19 plasmid DNA and observed >96.5%, suggestion false negative rate of less than 3.5%.

#### Hi-C data processing

The filtered fend-transformed read pairs were converted into “misha” tracks and imported into the genomic mm10 database. They were normalized using the Shaman package (https://tanaylab.bitbucket.io/shaman/index.html) and the hi-c score was calculated using a kNN strategy on the pooled replicates as previously described (Bonev et al., 2017) with a kNN of 100.

#### Contact probability, insulation and TAD boundary calling

Contact probability as a function of the genomic distance was calculated as previously described (Bonev et al., 2017). The data was presented as a log10 contact probability in log10 genomic distance bins. To define insulation based on observed contacts we used the insulation score as previously defined (Bonev et al., 2017; Nagano et al., 2017). The insulation score was computed on the pooled contact map at 1 kb resolution within a region of ±250 kb and is multiplied by (−1) so that high insulation score represents strong insulation. Domain boundaries were then defined as the local 2 kb maxima in regions, where the insulation score is above the 90% quantile of the genome-wide distribution. Differential TAD boundaries were identified as previously described (Bonev et al., 2017) using genome-wide normalized insulation scores.

#### Compartments and compartment strength

We first calculated the dominant eigenvector of the contact matrices binned at 250 kb as described (Lieberman-Aiden et al., 2009) using scripts available at (https://github.com/dekkerlab/cworld-dekker). To determine the compartment strength, we plotted the log2 ratio of observed versus expected contacts (intrachromosomal separated by at least 10Mb) either between domains of the same (A-A, B-B) or different type (A-B), as previously described (Bonev et al., 2017). We calculated compartment strength as the ratio between the sum of observed contacts within the A and B compartment and the sum of intercompartment contacts (AA+BB)/(AB+BA).

#### Average TAD contact enrichment

The insulation and contact enrichment within TADs was calculated as previously described (Bonev et al., 2017). Briefly, TAD coordinates were extended upstream and downstream by the TAD length and this distance was split into 100 equal bins. The observed vs expected enrichment ratio was calculated in each of the resulting 100×100 grid (per TAD) and the average enrichment was plotted per bin. Average CpG DNA methylation was calculated for each of these 100 bins per TAD and was represented as mean ± 0.25 quantiles.

#### Aggregated and individual contact strength at pairs of genomic features

To calculate the contact enrichment ratio at pairs of genomic features (such as ChIP-seq peaks, accessible motif sites or linked pairs of enhancers and promoters), we used two complementary approaches. First, we aggregated Hi-C maps to calculate the log2 ratio of the observed vs expected contacts within a window of a specific size, centred on the pair of interest, as described previously (Bonev et al., 2017). Furthermore, we calculated the average enrichment ratio of the contact strength in the center of the window (central 9 bins) vs each of the corners. This kind of analysis is useful to identify general patterns of changes in chromatin interactions in the data, but cannot distinguish the heterogeneity and the contribution of individual pairs. To address this question, we also extracted the kNN-based Hi-C score in a 10kb window centered around each of the pairs separately and represented the data as a scatter plot or boxplot. Significance was then calculated using Wilcoxon rank test.

#### Average enrichment of linear marks at genomic features

We used SeqPlots (Stempor and Ahringer, 2016) to calculate the average enrichment of linear chromatin marks (DNA methylation, chromatin accessibility, ChIP-seq) in window centered around the genomic feature of interest, or along scaled gene bodies.

#### Inferring Hi-C contact strength and CpG DNA methylation associated with TF binding motifs

Although ChIP-seq (and related techniques) remains the method of choice to identify the real binding sites of a TF, in many cases this is not feasible due to the lack of suitable antibodies. We reasoned that we could use predicted TF motifs at highly accessible peaks to examine how such potentially occupied motifs are spatially positioned to each other and how they are associated with changes in DNA methylation. Briefly, for each transcription factor motif we first identified the scATAC-based peaks which contain the predicted binding motif and then ranked these cites based on their maximum accessibility in aggregated pseudobulk scATAC-based clusters. To not overpenalize rare motifs, we selected the top 5000 most highly accessible regions per motif and created point-based regions centered at the corresponding motif. TFs (and their corresponding motifs) which are not expressed in our data (RPKM>1 based on pseudobulk scRNA-seq data) or are not within the top 3000 most variable genes (based on scRNA-seq) were discarded from further analysis.

To create a set of control regions, not enriched for any particular motif, we first selected a set of the top 50000 most highly accessible sites, analogous to motif-based analysis. Then, we sampled 5000 peaks randomly 1000 times to create a set of highly robust background peaks with similar characteristics. Next, we created pairs of these regions separated by at least 10kb, filtered for intra-TAD interactions and extracted the maximum Hi-C score (for each cell type) within a square window of 10×10 kb centered on each pair. We then calculated the median value per transcription factor motif per cell type and repeated this for each of the 1000 controls to generate a background distribution. To normalize for any non-specific interaction, we calculated the mean of the 1000 controls per cell type and subtracted it from the TF motif values. We then calculated the maximum Hi-C score (from all the three cell type) and the standard deviation per motif and plotted them as a scatter plot using ggplot2. To calculate the statistical significance, we used permutation based analysis (utilizing the 1000 repeated sampling of the random regions) and considered motifs with p<0.05 as significant. We also depicted the Pearson’s correlation between the motif deviation scores and the expression of the matching TF as described for Figure 3E.

For the analysis of DNA methylation, we extracted the average CpG methylation in 500 bp windows centered on the motif-based regions described above. We then generated a random permutation-based background exactly as described for the Hi-C and plotted the minimum DNA methylation and standard deviation for the three cell types as a scatter plot. Statistical significance was calculated as described above.

As a complimentary analysis we calculated the average Hi-C score and enhancer DNA methylation for positively correlated enhancer-gene pairs and presented the data as scaled median Hi-C score per TF motif.

#### Software

Methyl-Hi-C data was processed using the JuiceMe pipeline, available at https://github.com/aidenlab/JuiceMe. The R package to compute the expected tracks and the Hi-C scores is available at https://bitbucket.org/tanaylab/shaman.

#### Data Resources

Data will be deposited on GEO and made available upon publication.

## References

Aprea, J., Prenninger, S., Dori, M., Ghosh, T., Monasor, L.S., Wessendorf, E., Zocher, S., Massalini, S., Alexopoulou, D., Lesche, M., et al. (2013). Transcriptome sequencing during mouse brain development identifies long non-coding RNAs functionally involved in neurogenic commitment. EMBO J. 32, 3145–3160.

Argelaguet, R., Clark, S.J., Mohammed, H., Stapel, L.C., Krueger, C., Kapourani, C.-A., Imaz-Rosshandler, I., Lohoff, T., Xiang, Y., Hanna, C.W., et al. (2019). Multi-omics profiling of mouse gastrulation at single-cell resolution. Nature 576, 487–491.

Argelaguet, R., Arnol, D., Bredikhin, D., Deloro, Y., Velten, B., Marioni, J.C., and Stegle, O. (2020). MOFA+: a statistical framework for comprehensive integration of multi-modal single-cell data. Genome Biol. 21, 111.

Bayam, E., Sahin, G.S., Guzelsoy, G., Guner, G., Kabakcioglu, A., and Ince-Dunn, G. (2015). Genome-wide target analysis of NEUROD2 provides new insights into regulation of cortical projection neuron migration and differentiation. BMC Genomics 16, 681.

Berest, I., Arnold, C., Reyes-Palomares, A., Palla, G., Rasmussen, K.D., Giles, H., Bruch, P.-M., Huber, W., Dietrich, S., Helin, K., et al. (2019). Quantification of Differential Transcription Factor Activity and Multiomics-Based Classification into Activators and Repressors: diffTF. Cell Rep. 29, 3147-3159.e12.

Bergen, V., Lange, M., Peidli, S., Wolf, F.A., and Theis, F.J. (2020). Generalizing RNA velocity to transient cell states through dynamical modeling. Nat. Biotechnol. 1–7.

Bonev, B., and Cavalli, G. (2016). Organization and function of the 3D genome. Nat Rev Genet 17, 661–678.

Bonev, B., Mendelson Cohen, N., Szabo, Q., Fritsch, L., Papadopoulos, G.L., Lubling, Y., Xu, X., Lv, X., Hugnot, J.-P., Tanay, A., et al. (2017). Multiscale 3D Genome Rewiring during Mouse Neural Development. Cell 171, 557-572.e24.

de Bruijn, D.R.H., van Dijk, A.H.A., Pfundt, R., Hoischen, A., Merkx, G.F.M., Gradek, G.A., Lybæk, H., Stray-Pedersen, A., Brunner, H.G., and Houge, G. (2010). Severe Progressive Autism Associated with Two de novo Changes: A 2.6-Mb 2q31.1 Deletion and a Balanced t(14;21)(q21.1;p11.2) Translocation with Long-Range Epigenetic Silencing of LRFN5 Expression. Mol. Syndromol. 1, 46–57.

Cao, J., Spielmann, M., Qiu, X., Huang, X., Ibrahim, D.M., Hill, A.J., Zhang, F., Mundlos, S., Christiansen, L., Steemers, F.J., et al. (2019). The single-cell transcriptional landscape of mammalian organogenesis. Nature 566, 496–502.

Choi, Y., Nam, J., Whitcomb, D.J., Song, Y.S., Kim, D., Jeon, S., Um, J.W., Lee, S.-G., Woo, J., Kwon, S.-K., et al. (2016). SALM5 trans-synaptically interacts with LAR-RPTPs in a splicing-dependent manner to regulate synapse development. Sci. Rep. 6, 26676.

Clark, S.J., Argelaguet, R., Kapourani, C.-A., Stubbs, T.M., Lee, H.J., Alda-Catalinas, C., Krueger, F., Sanguinetti, G., Kelsey, G., Marioni, J.C., et al. (2018). scNMT-seq enables joint profiling of chromatin accessibility DNA methylation and transcription in single cells. Nat Commun 9, 781.

Corces, M.R., Buenrostro, J.D., Wu, B., Greenside, P.G., Chan, S.M., Koenig, J.L., Snyder, M.P., Pritchard, J.K., Kundaje, A., Greenleaf, W.J., et al. (2016). Lineage-specific and single cell chromatin accessibility charts human hematopoiesis and leukemia evolution. Nat. Genet. 48, 1193–1203.

Corces, M.R., Granja, J.M., Shams, S., Louie, B.H., Seoane, J.A., Zhou, W., Silva, T.C., Groeneveld, C., Wong, C.K., Cho, S.W., et al. (2018). The chromatin accessibility landscape of primary human cancers. Science 362, eaav1898.

Cowling, B.S., Cottle, D.L., Wilding, B.R., D’Arcy, C.E., Mitchell, C.A., and McGrath, M.J. (2011). Four and a half LIM protein 1 gene mutations cause four distinct human myopathies: a comprehensive review of the clinical, histological and pathological features. Neuromuscul. Disord. NMD 21, 237–251.

Deng, W., Lee, J., Wang, H., Miller, J., Reik, A., Gregory, P.D., Dean, A., and Blobel, G.A. (2012). Controlling long-range genomic interactions at a native locus by targeted tethering of a looping factor. Cell 149, 1233–1244.

Domcke, S., Bardet, A.F., Adrian Ginno, P., Hartl, D., Burger, L., and Schübeler, D. (2015). Competition between DNA methylation and transcription factors determines binding of NRF1. Nature 528, 575–579.

Durand, N.C., Shamim, M.S., Machol, I., Rao, S.S.P., Huntley, M.H., Lander, E.S., and Aiden, E.L. (2016). Juicer Provides a One-Click System for Analyzing Loop-Resolution Hi-C Experiments. Cell Syst. 3, 95–98.

Estève, P.-O., Vishnu, U.S., Chin, H.G., and Pradhan, S. (2020). Visualization and sequencing of accessible chromatin reveals cell cycle and post romidepsin treatment dynamics. BioRxiv 2020.04.27.064691.

Farkas, L.M., Haffner, C., Giger, T., Khaitovich, P., Nowick, K., Birchmeier, C., Pääbo, S., and Huttner, W.B. (2008). Insulinoma-Associated 1 Has a Panneurogenic Role and Promotes the Generation and Expansion of Basal Progenitors in the Developing Mouse Neocortex. Neuron 60, 40–55.

Feng, C., Song, C., Liu, Y., Qian, F., Gao, Y., Ning, Z., Wang, Q., Jiang, Y., Li, Y., Li, M., et al. (2020). KnockTF: a comprehensive human gene expression profile database with knockdown/knockout of transcription factors. Nucleic Acids Res. 48, D93–D100.

Feng, W., Khan, M.A., Bellvis, P., Zhu, Z., Bernhardt, O., Herold-Mende, C., and Liu, H.-K. (2013). The chromatin remodeler CHD7 regulates adult neurogenesis via activation of SoxC transcription factors. Cell Stem Cell 13, 62–72.

Feng, W., Kawauchi, D., Körkel-Qu, H., Deng, H., Serger, E., Sieber, L., Lieberman, J.A., Jimeno-González, S., Lambo, S., Hanna, B.S., et al. (2017). Chd7 is indispensable for mammalian brain development through activation of a neuronal differentiation programme. Nat. Commun. 8, 14758.

Finak, G., McDavid, A., Yajima, M., Deng, J., Gersuk, V., Shalek, A.K., Slichter, C.K., Miller, H.W., McElrath, M.J., Prlic, M., et al. (2015). MAST: a flexible statistical framework for assessing transcriptional changes and characterizing heterogeneity in single-cell RNA sequencing data. Genome Biol. 16, 278.

Fornes, O., Castro-Mondragon, J.A., Khan, A., van der Lee, R., Zhang, X., Richmond, P.A., Modi, B.P., Correard, S., Gheorghe, M., Baranašić, D., et al. (2020). JASPAR 2020: update of the open-access database of transcription factor binding profiles. Nucleic Acids Res. 48, D87–D92.

Ghavi-Helm, Y., Klein, F.A., Pakozdi, T., Ciglar, L., Noordermeer, D., Huber, W., and Furlong, E.E.M. (2014). Enhancer loops appear stable during development and are associated with paused polymerase. Nature 512, 96–100.

Ghavi-Helm, Y., Jankowski, A., Meiers, S., Viales, R.R., Korbel, J.O., and Furlong, E.E.M. (2019). Highly rearranged chromosomes reveal uncoupling between genome topology and gene expression. Nat. Genet. 51, 1272–1282.

Gorkin, D.U., Barozzi, I., Zhao, Y., Zhang, Y., Huang, H., Lee, A.Y., Li, B., Chiou, J., Wildberg, A., Ding, B., et al. (2020). An atlas of dynamic chromatin landscapes in mouse fetal development. Nature 583, 744–751.

Götz, M., and Huttner, W.B. (2005). The cell biology of neurogenesis. Nat. Rev. Mol. Cell Biol. 6, 777–788.

Govindan, S., and Jabaudon, D. (2017). Coupling progenitor and neuronal diversity in the developing neocortex. FEBS Lett. 591, 3960–3977.

Granja, J.M., Klemm, S., McGinnis, L.M., Kathiria, A.S., Mezger, A., Corces, M.R., Parks, B., Gars, E., Liedtke, M., Zheng, G.X.Y., et al. (2019). Single-cell multiomic analysis identifies regulatory programs in mixed-phenotype acute leukemia. Nat. Biotechnol.

Granja, J.M., Corces, M.R., Pierce, S.E., Bagdatli, S.T., Choudhry, H., Chang, H.Y., and Greenleaf, W.J. (2020). ArchR: An integrative and scalable software package for single-cell chromatin accessibility analysis. BioRxiv 2020.04.28.066498.

Hahn, M.A., Jin, S.-G., Li, A.X., Liu, J., Huang, Z., Wu, X., Kim, B.-W., Johnson, J., Bilbao, A.-D.V., Tao, S., et al. (2019). Reprogramming of DNA methylation at NEUROD2-bound sequences during cortical neuron differentiation. Sci. Adv. 5, eaax0080.

He, P., Williams, B.A., Trout, D., Marinov, G.K., Amrhein, H., Berghella, L., Goh, S.-T., Plajzer-Frick, I., Afzal, V., Pennacchio, L.A., et al. (2020). The changing mouse embryo transcriptome at whole tissue and single-cell resolution. Nature 583, 760–767.

Heng, J.I.-T., Nguyen, L., Castro, D.S., Zimmer, C., Wildner, H., Armant, O., Skowronska-Krawczyk, D., Bedogni, F., Matter, J.-M., Hevner, R., et al. (2008). Neurogenin 2 controls cortical neuron migration through regulation of Rnd2. Nature 455, 114–118.

Hevner, R.F. (2019). Intermediate progenitors and Tbr2 in cortical development. J. Anat. 235, 616–625.

Hsiung, C.C.-S., Morrissey, C.S., Udugama, M., Frank, C.L., Keller, C.A., Baek, S., Giardine, B., Crawford, G.E., Sung, M.-H., Hardison, R.C., et al. (2015). Genome accessibility is widely preserved and locally modulated during mitosis. Genome Res. 25, 213–225.

Javierre, B.M., Burren, O.S., Wilder, S.P., Kreuzhuber, R., Hill, S.M., Sewitz, S., Cairns, J., Wingett, S.W., Várnai, C., Thiecke, M.J., et al. (2016). Lineage-Specific Genome Architecture Links Enhancers and Non-coding Disease Variants to Target Gene Promoters. Cell 167, 1369-1384.e19.

Klemm, S.L., Shipony, Z., and Greenleaf, W.J. (2019). Chromatin accessibility and the regulatory epigenome. Nat. Rev. Genet. 20, 207–220.

Korsunsky, I., Millard, N., Fan, J., Slowikowski, K., Zhang, F., Wei, K., Baglaenko, Y., Brenner, M., Loh, P., and Raychaudhuri, S. (2019). Fast, sensitive and accurate integration of single-cell data with Harmony. Nat. Methods 16, 1289–1296.

La Manno, G., Soldatov, R., Zeisel, A., Braun, E., Hochgerner, H., Petukhov, V., Lidschreiber, K., Kastriti, M.E., Lönnerberg, P., Furlan, A., et al. (2018). RNA velocity of single cells. Nature 560, 494–498.

Lee, D.-S., Luo, C., Zhou, J., Chandran, S., Rivkin, A., Bartlett, A., Nery, J.R., Fitzpatrick, C., O’Connor, C., Dixon, J.R., et al. (2019). Simultaneous profiling of 3D genome structure and DNA methylation in single human cells. Nat. Methods 16, 999–1006.

Li, G., Liu, Y., Zhang, Y., Kubo, N., Yu, M., Fang, R., Kellis, M., and Ren, B. (2019). Joint profiling of DNA methylation and chromatin architecture in single cells. Nat. Methods 16, 991–993.

Lieberman-Aiden, E., Berkum, N.L. van, Williams, L., Imakaev, M., Ragoczy, T., Telling, A., Amit, I., Lajoie, B.R., Sabo, P.J., Dorschner, M.O., et al. (2009). Comprehensive Mapping of Long-Range Interactions Reveals Folding Principles of the Human Genome. Science 326, 289–293.

Llinares-Benadero, C., and Borrell, V. (2019). Deconstructing cortical folding: genetic, cellular and mechanical determinants. Nat. Rev. Neurosci.

Loo, L., Simon, J.M., Xing, L., McCoy, E.S., Niehaus, J.K., Guo, J., Anton, E.S., and Zylka, M.J. (2019). Single-cell transcriptomic analysis of mouse neocortical development. Nat. Commun. 10, 1–11.

Lupiáñez, D.G., Kraft, K., Heinrich, V., Krawitz, P., Brancati, F., Klopocki, E., Horn, D., Kayserili, H., Opitz, J.M., Laxova, R., et al. (2015). Disruptions of topological chromatin domains cause pathogenic rewiring of gene-enhancer interactions. Cell 161, 1012–1025.

Ma, S., Zhang, B., LaFave, L., Chiang, Z., Hu, Y., Ding, J., Brack, A., Kartha, V.K., Law, T., Lareau, C., et al. (2020). Chromatin potential identified by shared single cell profiling of RNA and chromatin. BioRxiv 2020.06.17.156943.

Mattar, P., Langevin, L.M., Markham, K., Klenin, N., Shivji, S., Zinyk, D., and Schuurmans, C. (2008). Basic helix-loop-helix transcription factors cooperate to specify a cortical projection neuron identity. Mol. Cell. Biol. 28, 1456–1469.

Meissner, A., Mikkelsen, T.S., Gu, H., Wernig, M., Hanna, J., Sivachenko, A., Zhang, X., Bernstein, B.E., Nusbaum, C., Jaffe, D.B., et al. (2008). Genome-scale DNA methylation maps of pluripotent and differentiated cells. Nature 454, 766–770.

Mikhail, F.M., Lose, E.J., Robin, N.H., Descartes, M.D., Rutledge, K.D., Rutledge, S.L., Korf, B.R., and Carroll, A.J. (2011). Clinically relevant single gene or intragenic deletions encompassing critical neurodevelopmental genes in patients with developmental delay, mental retardation, and/or autism spectrum disorders. Am. J. Med. Genet. A. 155A, 2386–2396.

Monaghan, C.E., Nechiporuk, T., Jeng, S., McWeeney, S.K., Wang, J., Rosenfeld, M.G., and Mandel, G. (2017). REST corepressors RCOR1 and RCOR2 and the repressor INSM1 regulate the proliferation–differentiation balance in the developing brain. Proc. Natl. Acad. Sci. 114, E406–E415.

Mukhtar, T., Breda, J., Grison, A., Karimaddini, Z., Grobecker, P., Iber, D., Beisel, C., van Nimwegen, E., and Taylor, V. (2020). Tead transcription factors differentially regulate cortical development. Sci. Rep. 10.

Nagano, T., Lubling, Y., Várnai, C., Dudley, C., Leung, W., Baran, Y., Mendelson Cohen, N., Wingett, S., Fraser, P., and Tanay, A. (2017). Cell-cycle dynamics of chromosomal organization at single-cell resolution. Nature 547, 61–67.

Noack, F., Pataskar, A., Schneider, M., Buchholz, F., Tiwari, V.K., and Calegari, F. (2019). Assessment and site-specific manipulation of DNA (hydroxy-)methylation during mouse corticogenesis. Life Sci. Alliance 2.

Nora, E.P., Goloborodko, A., Valton, A.-L., Gibcus, J.H., Uebersohn, A., Abdennur, N., Dekker, J., Mirny, L.A., and Bruneau, B.G. (2017). Targeted Degradation of CTCF Decouples Local Insulation of Chromosome Domains from Genomic Compartmentalization. Cell 169, 930-944.e22.

Pagin, M., Giubbolini, S., Barone, C., Sambruni, G., Zhu, Y., Ottolenghi, S., Wei, C.-L., and Nicolis, S.K. (2020). Sox2 controls neural stem cell self-renewal through a Fos-centered gene regulatory network. BioRxiv 2020.03.17.995621.

Pereira, J.D., Sansom, S.N., Smith, J., Dobenecker, M.-W., Tarakhovsky, A., and Livesey, F.J. (2010). Ezh2, the histone methyltransferase of PRC2, regulates the balance between self-renewal and differentiation in the cerebral cortex. Proc. Natl. Acad. Sci. 107, 15957–15962.

Pinto, L., Drechsel, D., Schmid, M.-T., Ninkovic, J., Irmler, M., Brill, M.S., Restani, L., Gianfranceschi, L., Cerri, C., Weber, S.N., et al. (2009). AP2gamma regulates basal progenitor fate in a region- and layer-specific manner in the developing cortex. Nat. Neurosci. 12, 1229–1237.

Preissl, S., Fang, R., Huang, H., Zhao, Y., Raviram, R., Gorkin, D.U., Zhang, Y., Sos, B.C., Afzal, V., Dickel, D.E., et al. (2018). Single-nucleus analysis of accessible chromatin in developing mouse forebrain reveals cell-type-specific transcriptional regulation. Nat. Neurosci. 21, 432–439.

Rao, S.S.P., Huntley, M.H., Durand, N.C., Stamenova, E.K., Bochkov, I.D., Robinson, J.T., Sanborn, A., Machol, I., Omer, A.D., Lander, E.S., et al. (2014). A three-dimensional map of the human genome at kilobase resolution reveals principles of chromatin looping. Cell 159, 1665–1680.

Rubin, A.J., Barajas, B.C., Furlan-Magaril, M., Lopez-Pajares, V., Mumbach, M.R., Howard, I., Kim, D.S., Boxer, L.D., Cairns, J., Spivakov, M., et al. (2017). Lineage-specific dynamic and pre-established enhancer-promoter contacts cooperate in terminal differentiation. Nat. Genet. 49, 1522–1528.

Schep, A.N., Wu, B., Buenrostro, J.D., and Greenleaf, W.J. (2017). chromVAR: inferring transcription-factor-associated accessibility from single-cell epigenomic data. Nat. Methods 14, 975–978.

Schoenfelder, S., and Fraser, P. (2019). Long-range enhancer-promoter contacts in gene expression control. Nat. Rev. Genet. 20, 437–455.

Schuurmans, C., Armant, O., Nieto, M., Stenman, J.M., Britz, O., Klenin, N., Brown, C., Langevin, L.-M., Seibt, J., Tang, H., et al. (2004). Sequential phases of cortical specification involve Neurogenin-dependent and -independent pathways. EMBO J. 23, 2892–2902.

Sessa, A., Mao, C.-A., Hadjantonakis, A.-K., Klein, W.H., and Broccoli, V. (2008). Tbr2 directs conversion of radial glia into basal precursors and guides neuronal amplification by indirect neurogenesis in the developing neocortex. Neuron 60, 56–69.

Sessa, A., Ciabatti, E., Drechsel, D., Massimino, L., Colasante, G., Giannelli, S., Satoh, T., Akira, S., Guillemot, F., and Broccoli, V. (2017). The Tbr2 Molecular Network Controls Cortical Neuronal Differentiation Through Complementary Genetic and Epigenetic Pathways. Cereb. Cortex 27, 3378–3396.

Stadler, M.B., Murr, R., Burger, L., Ivanek, R., Lienert, F., Schöler, A., Nimwegen, E. van, Wirbelauer, C., Oakeley, E.J., Gaidatzis, D., et al. (2011). DNA-binding factors shape the mouse methylome at distal regulatory regions. Nature 480, 490–495.

Stempor, P., and Ahringer, J. (2016). SeqPlots - Interactive software for exploratory data analyses, pattern discovery and visualization in genomics. Wellcome Open Res. 1, 14.

Stuart, T., Butler, A., Hoffman, P., Hafemeister, C., Papalexi, E., Mauck, W.M., Hao, Y., Stoeckius, M., Smibert, P., and Satija, R. (2019). Comprehensive Integration of Single-Cell Data. Cell 177, 1888-1902.e21.

Stumm, R.K., Zhou, C., Schulz, S., Endres, M., Kronenberg, G., Allen, J.P., Tulipano, G., and Höllt, V. (2004). Somatostatin Receptor 2 Is Activated in Cortical Neurons and Contributes to Neurodegeneration after Focal Ischemia. J. Neurosci. 24, 11404–11415.

Sun, J., Rockowitz, S., Xie, Q., Ashery-Padan, R., Zheng, D., and Cvekl, A. (2015). Identification of in vivo DNA-binding mechanisms of Pax6 and reconstruction of Pax6-dependent gene regulatory networks during forebrain and lens development. Nucleic Acids Res. 43, 6827–6846.

Tavano, S., Taverna, E., Kalebic, N., Haffner, C., Namba, T., Dahl, A., Wilsch-Bräuninger, M., Paridaen, J.T.M.L., and Huttner, W.B. (2018). Insm1 Induces Neural Progenitor Delamination in Developing Neocortex via Downregulation of the Adherens Junction Belt-Specific Protein Plekha7. Neuron 97, 1299-1314.e8.

Telley, L., Govindan, S., Prados, J., Stevant, I., Nef, S., Dermitzakis, E., Dayer, A., and Jabaudon, D. (2016). Sequential transcriptional waves direct the differentiation of newborn neurons in the mouse neocortex. Science 351, 1443–1446.

Telley, L., Agirman, G., Prados, J., Amberg, N., Fièvre, S., Oberst, P., Bartolini, G., Vitali, I., Cadilhac, C., Hippenmeyer, S., et al. (2019). Temporal patterning of apical progenitors and their daughter neurons in the developing neocortex. Science 364, eaav2522.

Visel, A., Taher, L., Girgis, H., May, D., Golonzhka, O., Hoch, R.V., McKinsey, G.L., Pattabiraman, K., Silberberg, S.N., Blow, M.J., et al. (2013). A High-Resolution Enhancer Atlas of the Developing Telencephalon. Cell 152, 895–908.

Vissers, L.E.L.M., van Ravenswaaij, C.M.A., Admiraal, R., Hurst, J.A., de Vries, B.B.A., Janssen, I.M., van der Vliet, W.A., Huys, E.H.L.P.G., de Jong, P.J., Hamel, B.C.J., et al. (2004). Mutations in a new member of the chromodomain gene family cause CHARGE syndrome. Nat. Genet. 36, 955–957.

Zeisel, A., Muñoz-Manchado, A.B., Codeluppi, S., Lönnerberg, P., Manno, G.L., Juréus, A., Marques, S., Munguba, H., He, L., Betsholtz, C., et al. (2015). Cell types in the mouse cortex and hippocampus revealed by single-cell RNA-seq. Science 347, 1138–1142.

Zeisel, A., Hochgerner, H., Lönnerberg, P., Johnsson, A., Memic, F., van der Zwan, J., Häring, M., Braun, E., Borm, L.E., La Manno, G., et al. (2018). Molecular Architecture of the Mouse Nervous System. Cell 174, 999-1014.e22.

Zhu, H., Wang, G., and Qian, J. (2016). Transcription factors as readers and effectors of DNA methylation. Nat. Rev. Genet. 17, 551–565.

Ziller, M.J., Edri, R., Yaffe, Y., Donaghey, J., Pop, R., Mallard, W., Issner, R., Gifford, C.A., Goren, A., Xing, J., et al. (2015). Dissecting neural differentiation regulatory networks through epigenetic footprinting. Nature 518, 355–359.

