## Supplementary Figures for "Multiomics analysis reveals extensive epigenome remodeling during cortical development"

### Supplementary Figure 1

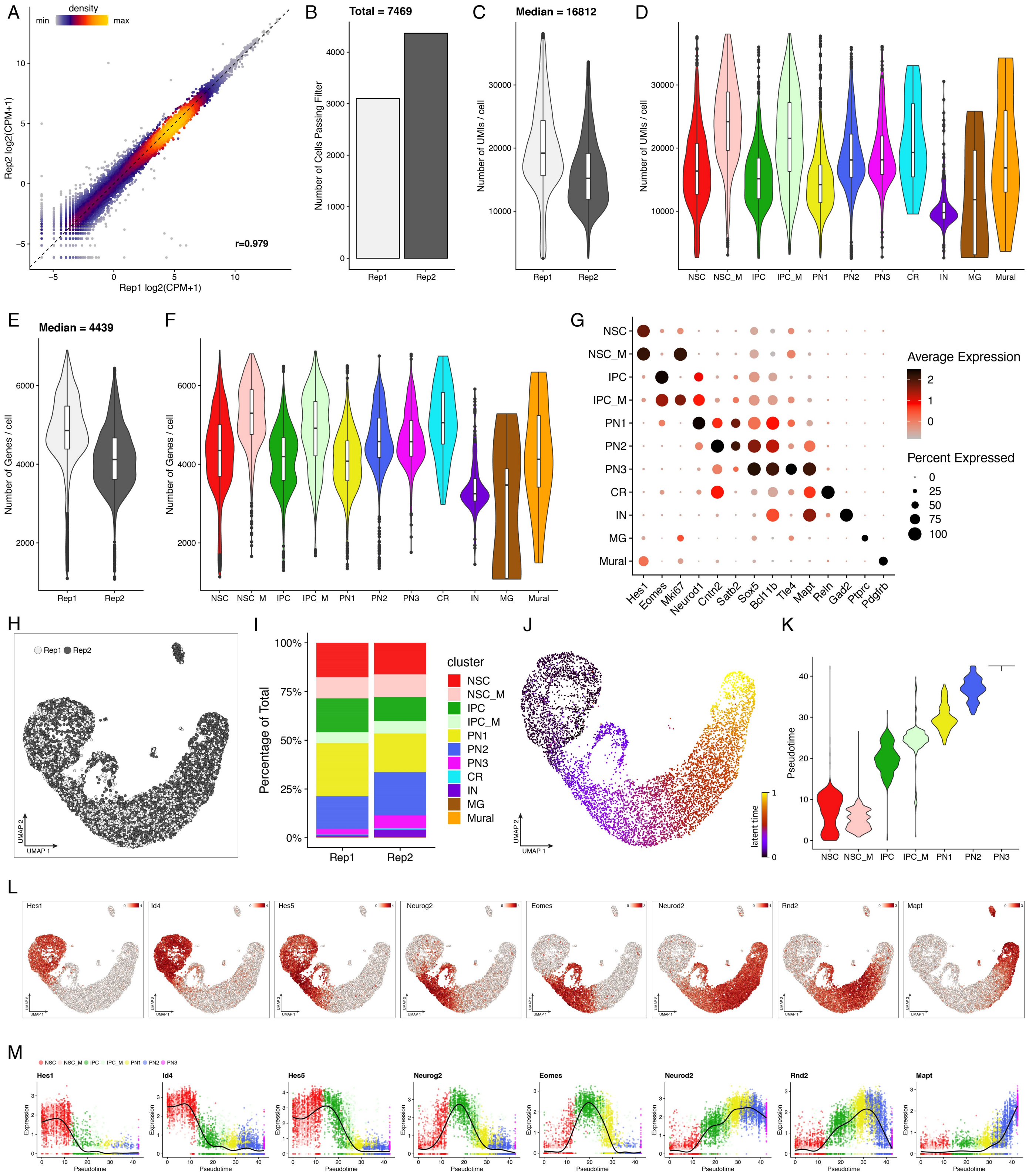


**Figure S1. Quality control and further validation of the scRNA-seq data**

(A) Scatterplot of gene expression values in the two biological scRNA-seq replicates, colored by density. The correlation (r) represents the Pearson correlation across all genes. (B) Barplot showing the number of cells passing quality filtering in each replicate. (C-D) Violin and box-whisker plots depicting the distribution of unique transcripts per cell across the two replicates and the identified clusters. (E-F) same as in (C-D) but showing the number of genes per cell. (G) Dot plot depicting the percentage of cells and gene expression levels of marker genes across the identified scRNA-seq clusters. (H) Overlay of replicate identity on the UMAP scRNA projection. (I) Stacked barplot showing the percentage of cells contributing to each cluster per replicate. (J) RNA velocity based inferred pseudotime depicted on the UMAP projection. Note the similarities with a trajectory-based approach (Figure 1F) (K) Violin plots depicting the distribution of pseudotime values per cluster. (L) UMAP visualization with expression levels of the indicated genes. (M) Gene expression changes along differentiation pseudotime. Each dot shows the expression per cell of the indicated gene, the line represents the smoothed fit of expression levels along pseudotime.

### Supplementary Figure 2

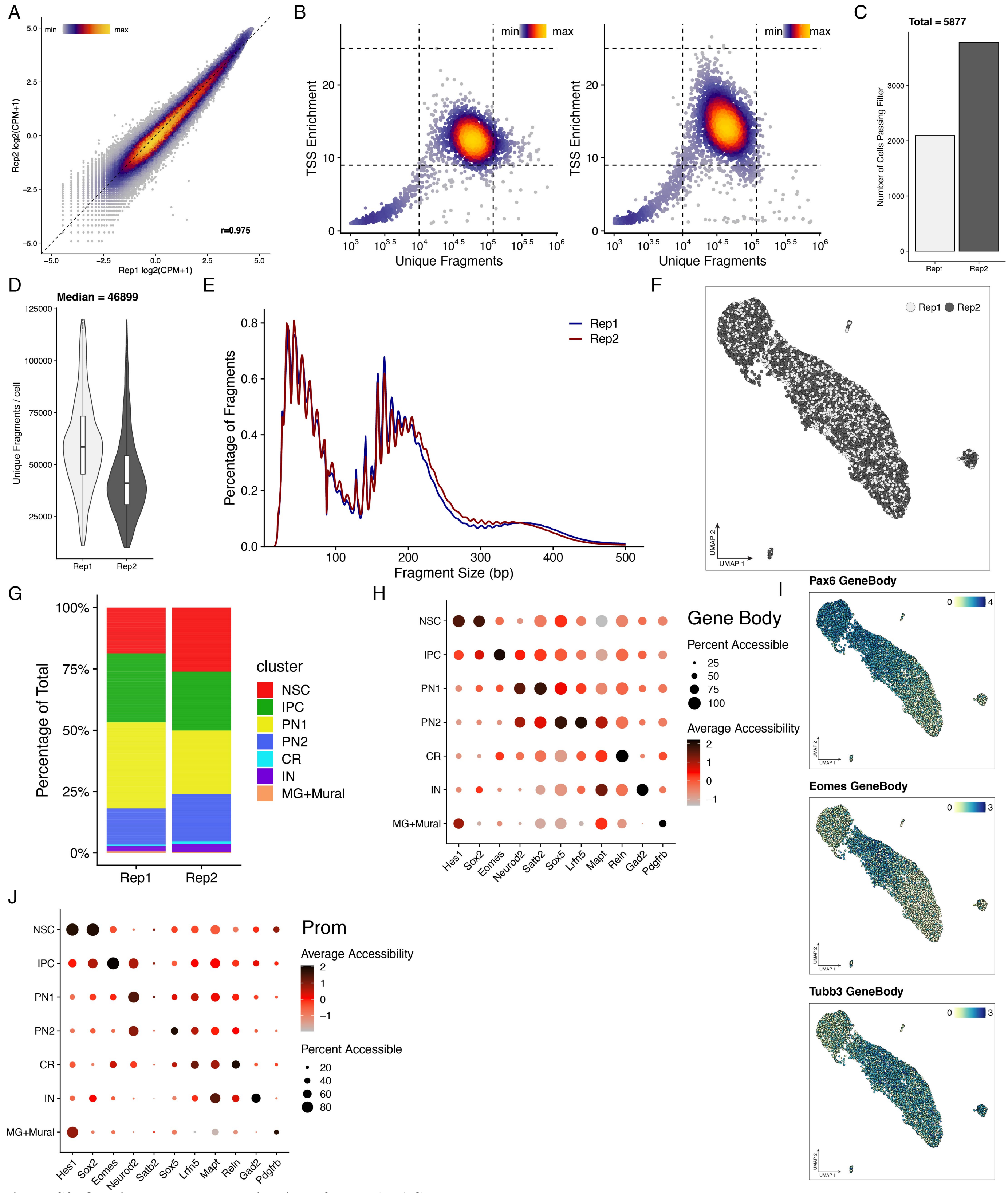


**Figure S2. Quality control and validation of the scATAC-seq data**

(A) Scatterplot of the aggregated accessibility values in the two biological scATAC-seq replicates, colored by density. The correlation (r) represents the Pearson correlation across all 370 975 peaks. (B) Scatter plot of the quality control metrics (TSS enrichment and unique fragments per cell) per replicate, colored by density. Dashed lines represent the filters for high-quality cells, retained for further analysis. (C) Barplot showing the number of cells passing quality filtering in each replicate. (D) Violin and box-whisker plots depicting the distribution of unique fragments per cell across the two replicates (E) Aggregated scATAC-seq fragment size distribution across replicates demonstrating sub-, mono- and multi-nucleosome spanning fragments. (F) Overlay of the replicate identity on the UMAP scATAC projection. (G) Stacked barplot showing the percentage of cells contributing to each cluster per replicate. (H) Dot plot depicting the percentage of cells and the average gene body accessibility of marker genes across the identified scATAC-seq clusters. (I) UMAP visualization with gene body accessibility of the indicated genes (J) Same as in (H) but based on promoter accessibility.

### Supplementary Figure 3

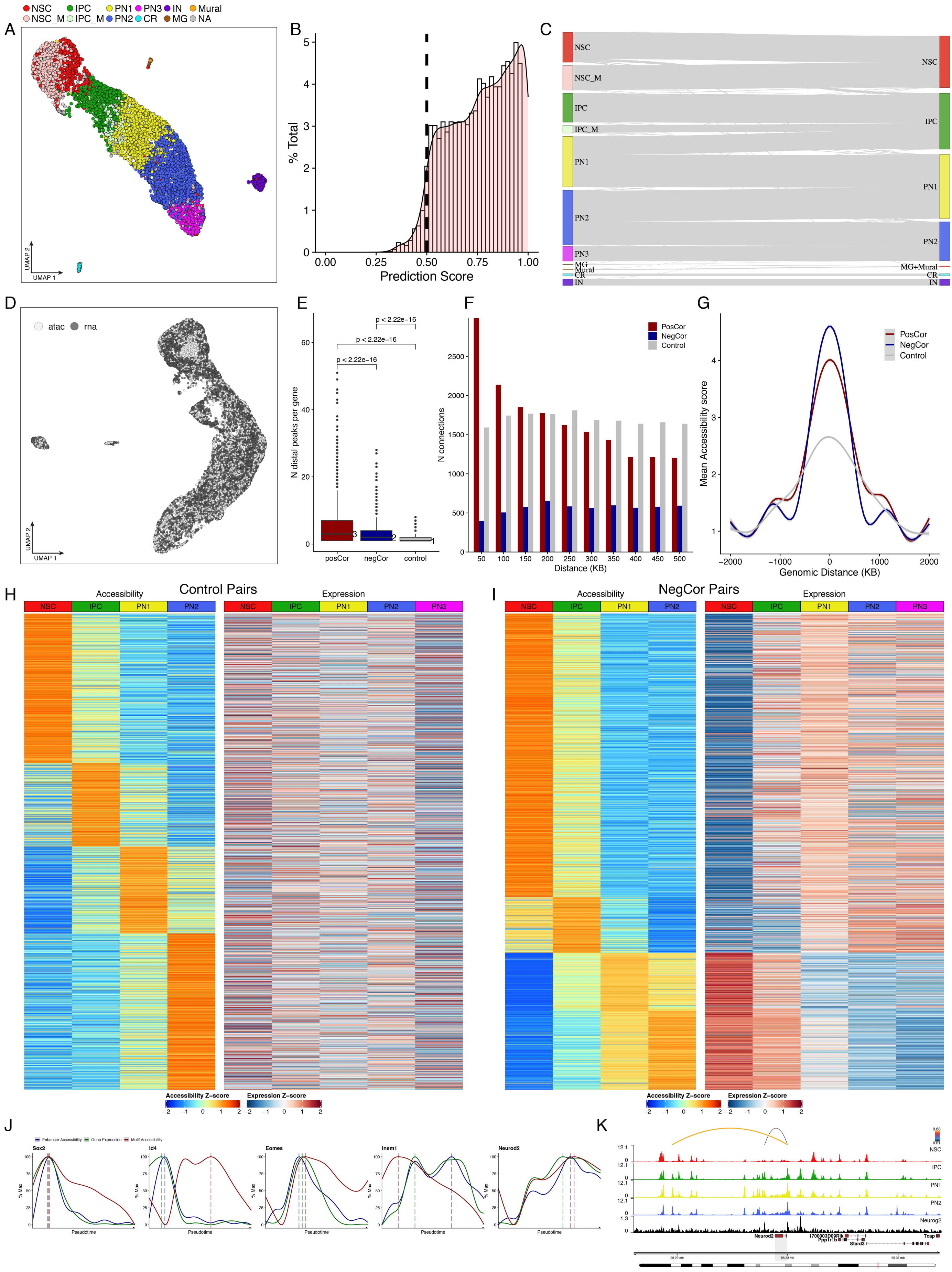

**Figure S3. scATAC-scRNA integration metrics and properties of the identified peak-gene links**

(A) scATAC UMAP projection with labels transferred from scRNA-seq. (B) Histogram-density plot depicting the distribution of prediction scores obtained during the scRNA-seq to scATAC-seq label transfer. Only cells with scores above the cutoff (dashed line) were retained. (C) Sankey plot showing the correspondence between scRNA-based labels (left) to scATAC-based cluster identities (right). (D) UMAP projection of the integrated scRNA-scATAC dataset, colored by method. (E) Box-whisker plot depicting the number of distal linked peaks per gene. Statistical significance is calculated using wilcoxon rank-sum test. Numbers represent median values per category. (F) Histogram displaying the genomic distance between TSS and positively, negatively or non-correlated (Control) peak-gene pairs. (G) Aggregated average accessibility plot (± 2kb) of the three categories of peak-gene pairs. (H-I) Heatmaps of aggregated accessibility of distal elements and gene expression levels of their linked genes for each of the identified non-correlated (Control) and negatively correlated pairs (NegCor). Rows were clustered by enhancer accessibility using feature binarization (Methods). (J) Smoothed fit line plots depicting the scaled aggregated enhancer accessibility, expression of the linked transcription factors and its motif accessibility across pseudotime. Dashed lines indicate the pseudo-temporal maxima. (K) Genomic tracks depicting the aggregated accessibility (per cluster) at the Neurod2 gene locus with arcs on top representing the identified enhancer-gene pairs, colored by Pearson correlation of the enhancer accessibility and gene expression. Neurog2 ChIPseq track is also depicted (black).

### Supplementary Figure 4

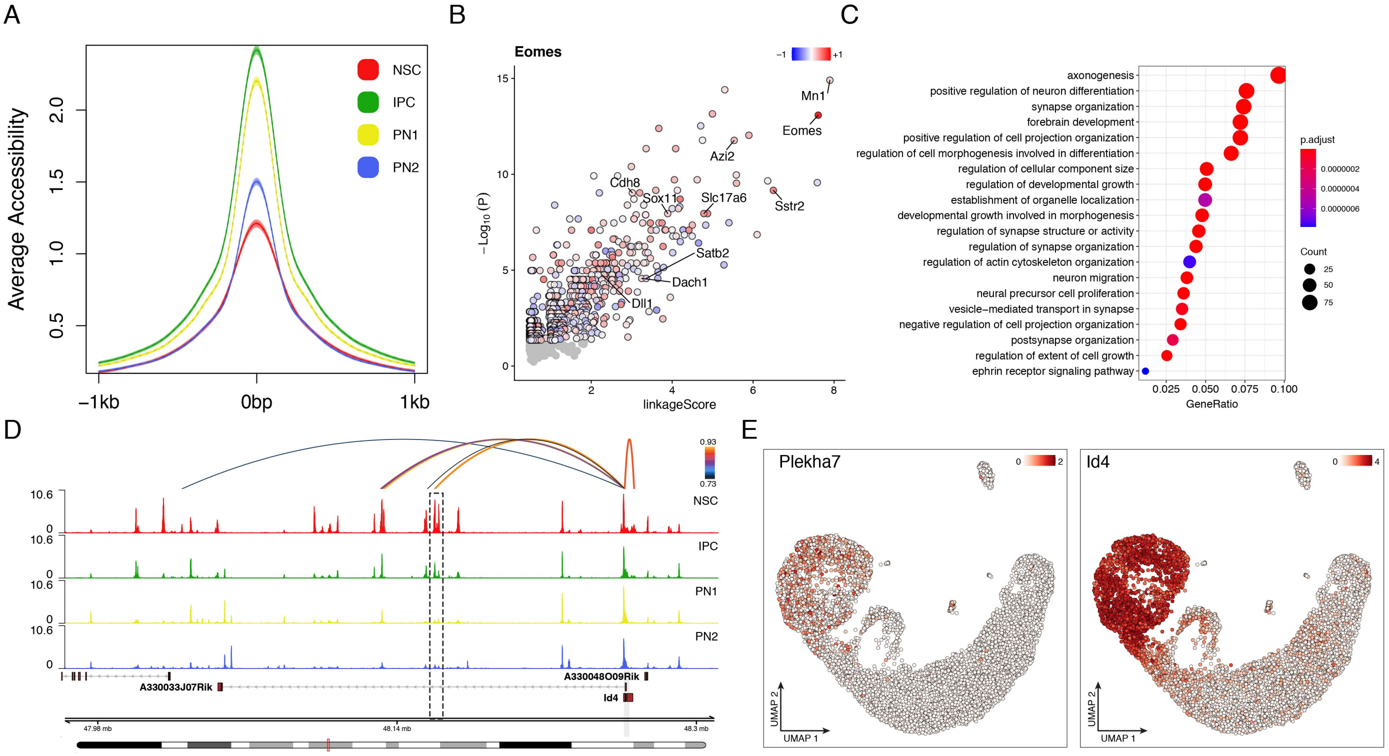


**Figure S4. Confirmation of the inferred gene regulatory networks for Eomes and Insm1**

(A) Average accessibility per cluster centered at ChIP-seq based Eomes binding sites (Sessa et al., 2017). (B) Scatter plot depicting the predicted downstream targets of Eomes based on the aggregated correlation between gene expression and accessibility of linked enhancers, overlapping with Eomes ChIP-seq peak (linkageScore) and peak enrichment. Genes are colored based on the Pearson correlation (r) between their expression and the expression of Eomes. Grey circles represent gene with non-significant peak enrichment (p>0.05; hypergeometric test). (C) Bar plot depicting the GO enriched terms, enrichment scores and gene ratios of the ChIP-seq based predicted Eomes targets. Note the high similarities of obtained GO terms from peak- and motif- (Figure 4C) centric analysis. (D) Genomic tracks depicting aggregated accessibility (per cluster) at the predicted Isnm1 target Id4. Arcs on top represent linked Id4 enhancers, overlapping with Insm1 motif and colored by Pearson correlation of the enhancer accessibility and Id4 expression. A VISTA-validated enhancer (Visel et al., 2013) is highlighted with a dashed rectangle. (E) scRNA UMAP projection, colored by the gene expression levels of Plekha7 (left) and Id4 (right).

#### Supplementary Figure 5

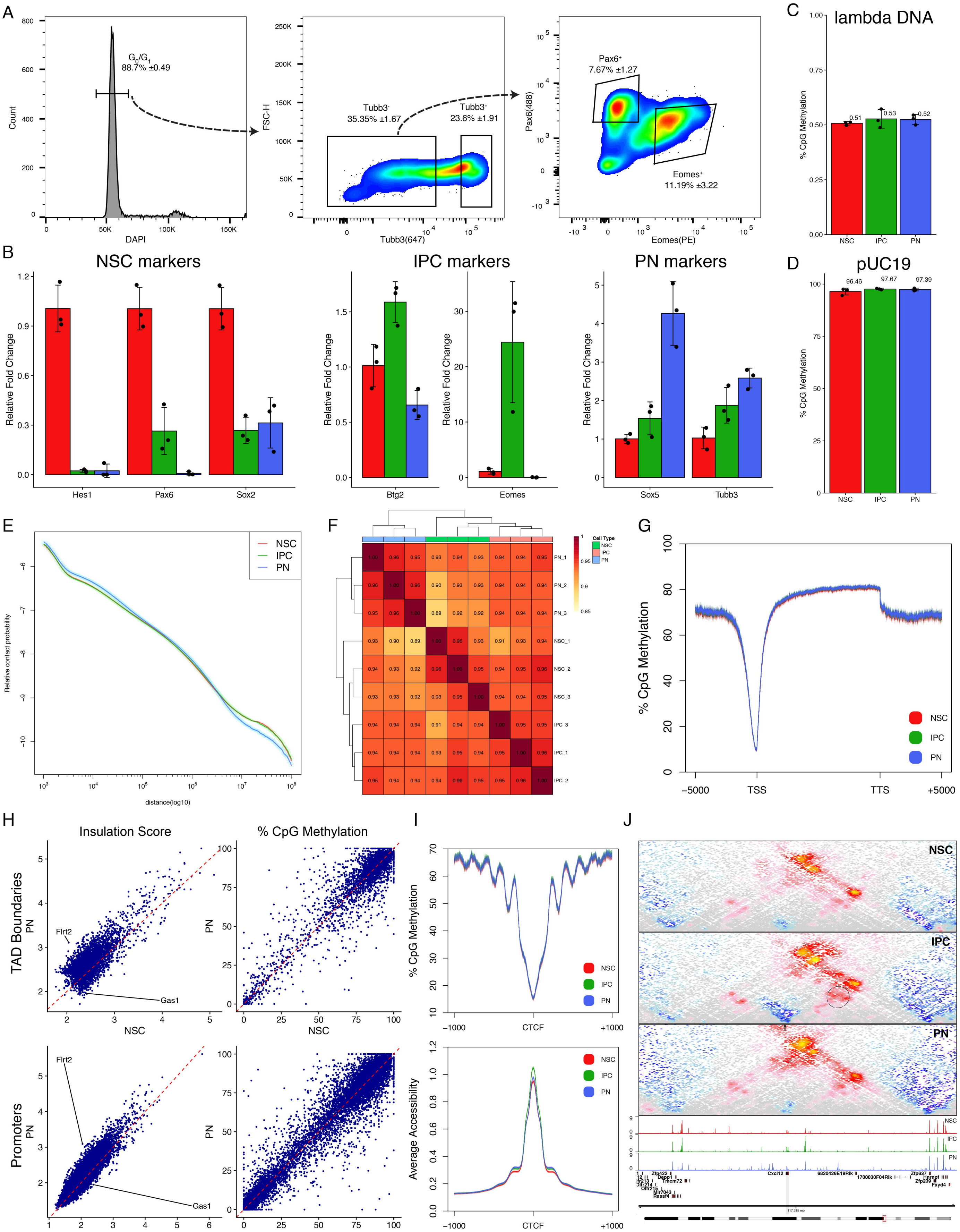

**Figure S5. Immuno-Methyl-Hi-C reveals global reorganization of the 3D genome architecture**
(A) Gating strategy for the immunoFAC-sorting of NSC, IP and PN. Singlets in $G_0/G_1$ based on their DNA content (DAPI) were selected first (left), followed by separation of nuclei with high or low Tubb3 signal (middle). Tubb3 low cells were further subdivided based on Pax6 and Eomes (right). Number represent mean ± SD from the parental singlet population. (B) Expression levels of NSC- (Hes1, Pax6, Sox2), IPC- (Btg2, Eomes) and PN- (Sox5, Tubb3) specific marker genes in the three isolated cell populations determined by RT-qPCR. Data are represented as a bar plot showing the mean ± SD, as well as individual biological replicates (black dots). Red= NSC, green= IPC, blue= PN (C-D) Bar plots depicting methylation levels (mean ± SD) of the spiked-in bisulfite conversion controls: either completely unmethylated lambda DNA (C), or fully methylated pUC19 DNA (D). Black dots indicate individual biological replicates. (E) Contact probability in logarithmic bins. Lines: mean values from biological replicates; semi-transparent ribbons: SEM (F) Pairwise Pearson's correlation between Hi-C samples (at 50 kb resolution and considering only contacts separated by at least 100 kb). Note that the major separation occurs between cell types. (G) Average DNA methylation across gene bodies (± 5 kb) in the indicated cell types. Note the stereotypical drop at DNA methylation at the TSS for all three cell types. (H) Scatterplot of insulation scores (left) and average DNA methylation levels (right) at all TAD boundaries (top) and gene promoters (bottom) in the indicated cell types. (I) Average DNA methylation levels of the three isolated cell types (top) and average accessibility from matched scATAC-seq clusters (bottom) at CTCF peaks (data from Bonev et al., 2017, NPC CTCF track). Note the nucleosomal pattern of DNA methylation around the CTCF sites. Lines: mean values from biological replicates; semi-transparent ribbons: SEM (J) Contact maps (top) and aggregated accessibility of matched scATAC-seq clusters (bottom) for a representative example of an IPC specific TAD boundary (arrow) at the Cxcl12 gene locus. Dynamic contacts are highlighted with a dashed ellipse.

### Supplementary Figure 6

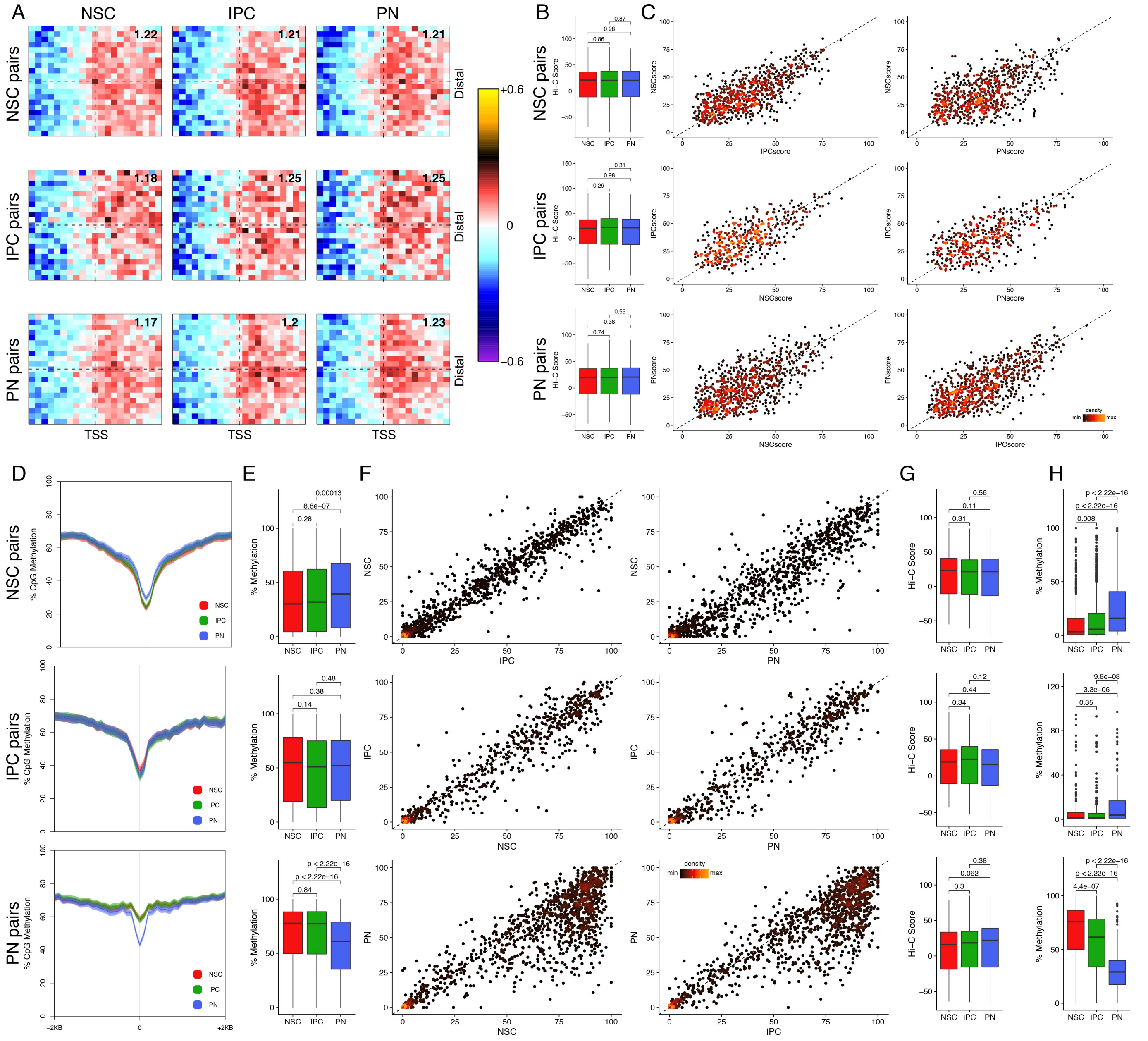


**Figure S6. Non-correlated enhancer-gene pairs are not associated with dynamic chromatin looping or changes at DNA methylation levels**
(A) Aggregated Hi-C maps between the linked distal peak and the transcription start site (TSS) of NSC-, IPC- and PN-specific non-correlated (Control) enhancer-gene pairs. Genes are oriented according to transcription. Number in the top-right corner indicates the ratio of the center enrichment to the mean of the four corners (Methods). (B-C) Box-whisker (B) and scatter plots colored by density (C), depicting the Hi-C score at non-correlated (Control) enhancer-gene pairs at the indicated cluster-specific pairs. Statistical significance is calculated using wilcoxon rank-sum test. (D) Average DNA methylation levels at the enhancers (± 2kb) of non-correlated (Control) correlated cluster-specific pairs. Lines: mean values from biological replicates; semi-transparent ribbons: SEM (E-F) Box-whisker (E) and scatter plots colored by density (F), depicting the average DNA methylation levels at enhancers of non-correlated (Control) cluster-specific pairs. Statistical significance is calculated using wilcoxon rank-sum test. (G-H) Box-whisker plots depicting the Hi-C scores at peak-gene pairs (G) or DNA methylation levels (H) at enhancers of negative correlated cluster-specific pairs.

### Supplementary Figure 7

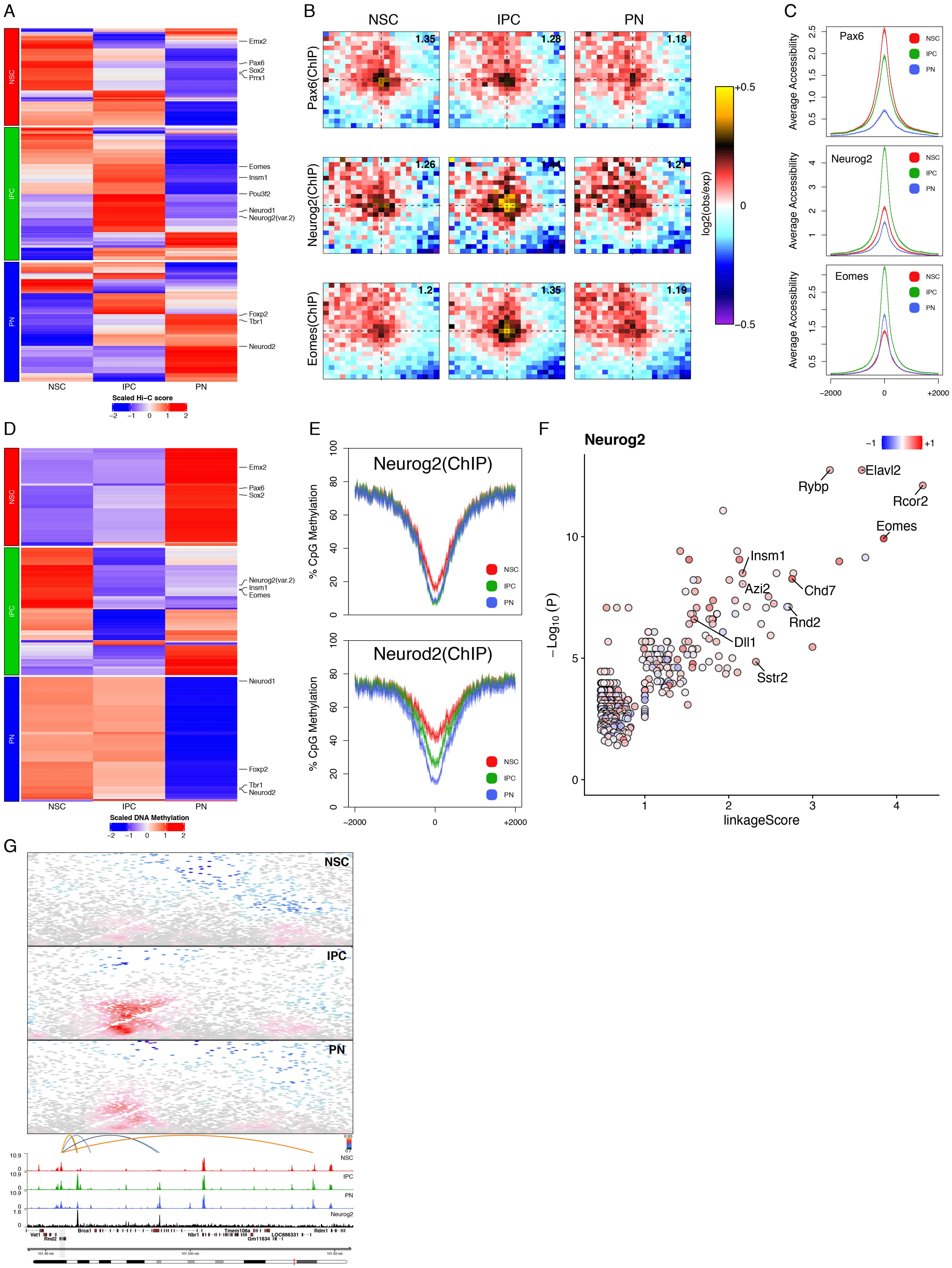

**Figure S7. Transcription factors associated with cell-type-specific looping based on positively correlated enhancer-gene pairs**

(A) Heatmaps depicting the scaled Hi-C score at positively correlated cluster-specific enhancer-gene pairs, where the TF motifs overlap with both the enhancer and the promoter. Only significant ($p<0.05$, permutation test) and expressed TFs are displayed. (B) Aggregated Hi-C plots of intraTAD pairs of ChIP-seq based bound sites. Number in the top-right corner indicates the ratio of the center enrichment to the mean of the four corners (Methods). (C) Average accessibility per cluster centered at the indicated ChIP-seq based binding sites (Sessa et al., 2017; Sun et al., 2015). (D) As (A) but plotting the scaled DNA methylation levels at enhancers, overlapping with TF motifs. (E) Average DNA methylation levels centered at ChIP-seq based Neurog2 (Sessa et al., 2017) and Neurod2 binding sites (Bayam et al., 2015). (F) Scatter plot depicting the predicted downstream targets of Neurog2 based on the aggregated correlation between gene expression and accessibility of linked enhancers, overlapping with Neurog2 ChIP-seq peak (linkageScore) and peak enrichment. Genes are colored based on the Pearson correlation between their expression and the expression of Neurog2. Grey circles represent gene with non-significant peak enrichment ($p>0.05$; hypergeometric test). (G) Contact maps (top) and aggregated accessibility of matched scATAC-seq clusters (bottom) at the Rnd2 locus. Arcs represent positively correlated enhancers-Rnd2 pairs and are colored based on the correlation between the enhancer accessibility and Rnd2 expression. Shown is also Neurog2 ChIP-seq track. Note the IPC-specific chromatin loops established between Neurog2-bound enhancers and Rnd2 promoter.
